# The development of slow and fast theta oscillations in the human brain

**DOI:** 10.64898/2026.09.27.754757

**Authors:** Zachariah R. Cross, Samantha M. Gray, Adam J. O. Dede, Yessenia M. Rivera, Qin Yin, Parisa Vahidi, Elias M. B. Rau, Christopher Cyr, Ania M. Holubecki, Eishi Asano, Olivia Kim McManus, Shifteh Sattar, Ignacio Saez, Fady Girgis, David King-Stephens, Peter B. Weber, Kenneth D. Laxer, Stephan U. Schuele, Joyce Y. Wu, Sandi K. Lam, Jeffrey S. Raskin, Ammar Shaikhouni, Peter Brunner, Jarod L. Roland, Rodrigo M. Braga, Robert T. Knight, Noa Ofen, Elizabeth L. Johnson

## Abstract

Theta oscillations are pivotal for attention and memory, with developmental shifts in theta activity and connectivity tracking memory ability across the lifespan. Building on initial evidence that the theta band separates into slow and fast sub-bands across development within the medial temporal lobe (MTL) and the prefrontal cortex (PFC), this preregistered study (https://osf.io/gsru7) characterized the frequencies of slow (∼1.5–4.5 Hz) and fast (∼4.5–9.0 Hz) theta oscillations across the brain in a large developmental cohort. We analyzed intracranial EEG (iEEG) recordings from widespread brain regions in 101 children and adults (5.93-54.00 years, 63 males; *n* electrodes = 5691) using task-based (attention to to-be-remembered visual stimuli) and task-free (resting-state) data. We reveal distinct slow and fast theta oscillations in all regions and identify a spatial gradient in both frequency ranges, with slow theta oscillations speeding up and fast theta oscillations slowing down from posterior to anterior regions. The dissociation between slow and fast theta frequencies peaks during young adulthood, mirroring the developmental trajectory of attention and memory. In the hippocampus and PFC, attentional state (task-based, task-free) modulated age effects, and individual differences in task-based slow theta frequencies predicted individual differences in memory performance, linking the development of slow theta oscillations to the development of memory. Gray matter volume was not associated with age- or task-related differences in theta frequencies, suggesting that the development of theta oscillations is independent of regional brain structure. This study establishes developmental trajectories of slow and fast theta oscillations in localized brain regions and links the maturation of hippocampal and PFC slow theta oscillations to the development of attention and memory.

## Introduction

Theta oscillations (∼2–9 Hz) dominate the medial temporal lobe (MTL) during memory encoding and retrieval processes ^1^, which undergo remarkable change across the lifespan ^2^, yet how theta oscillations develop remains poorly understood ^2,3^. Theta oscillations underpin information integration in the MTL ^4^ and synchronize activity between the MTL and regions such as the prefrontal cortex (PFC), supporting intra- and inter-regional coordination ^5–9^. While the developmental trajectory of alpha oscillations is well-documented ^10,11^, the developmental trajectory of theta oscillations, including how this coincides with shifts in cognitive abilities and brain structure has not been characterized. Understanding oscillatory theta dynamics across development is essential for understanding brain maturation and cognitive performance across the lifespan, foundational to basic and translational neuroscience.

Research on developmental differences in neural oscillations has traditionally relied on scalp-EEG (cf. ^9,12,13^), which cannot access medial regions, such as the MTL, and provides low spatial resolution ^3,14,15^. To address these limitations, this preregistered study (https://osf.io/gsru7) analyzed rare intracranial EEG (iEEG) data from an exceptionally large cohort of neurosurgical patients aged 5 to 54 years undergoing invasive monitoring for seizure management. In contrast to noninvasive neuroimaging, iEEG provides unparalleled access to the human brain, with millisecond temporal and millimeter spatial resolution, and a signal-to-noise ratio comparable to animal neurophysiology ^3,16–18^. After removing seizure zones and episodes of spread of seizure activity, iEEG data represents healthy neural activity ^19,20^, allowing a comprehensive investigation of localized theta oscillations across development.

Converging evidence from iEEG studies in humans and animal models suggests that theta oscillations in the MTL are crucial for binding disparate elements of experiences into coherent memory representations (for reviews, see ^4,21^). Coordinated theta oscillations between the MTL and the PFC are also associated with the orchestration of several cognitive processes, including episodic memory ^9,22^, decision-making ^23,24^, working memory ^25,26^, and spatial navigation ^27,28^. By modulating neuronal firing timing, theta oscillations enable the integration of information across networks ^25^, enhancing cognitive flexibility and adaptive neuronal functioning. These coordinated oscillations play a vital role in memory processing and the development of complex cognitive abilities ^29–32^.

Recent iEEG work revealed that distinct slow (∼1.5 – 4.5 Hz) and fast (∼4.5 – 9.0 Hz) theta oscillations in the MTL (here, parahippocampal gyrus) and the PFC undergo significant development from childhood to early adulthood ^9^. Across these regions, faster theta oscillations accelerate with age while slower theta oscillations decelerate, resulting in the separation of theta oscillations into two sub-bands as the brain matures into adulthood. However, these age-related differences in theta oscillations did not significantly explain developmental improvements in memory, suggesting that separation of slow and fast theta frequencies is intrinsic to brain maturation. Based on observing significant effects such as relationships between MTL-PFC coupling at individually-defined slow and fast theta frequencies and memory outcomes, the authors proposed that the separation of slow and fast theta oscillations might be indirectly linked to the maturation of memory systems as neural oscillations become more precise and differentiated ^2^. Yet, direct evidence is lacking. Critically, testing whether age-related differences in theta frequencies are themselves associated with memory performance in large, well-powered samples is a necessary first step toward testing this proposed pathway. Examining whole-brain theta frequencies will also determine whether regions outside of the MTL and PFC show age- and task-related differences that support memory development.

In this study, we defined regionally precise, brain-wide developmental trajectories of slow and fast theta frequencies in task-based and task-free states (Figures 1A, 1B). By comparing task-based and task-free frequencies, our approach revealed how localized brain regions, such as regions of the MTL and PFC, undergo functional specialization in coordinating memory and attention ^33^, linking oscillatory frequencies to cognitive development. In addition, we examined the relationship between regionally precise theta frequencies and cortical structure (Figure 1C). Measures of regional gray matter volume (GMV) and electrophysiological activity show substantial overlap in relation to cognition and age ^34–37^, which suggests that they may be jointly explained by shared factors, such as myelination and synaptogenesis. Thus, examining structure-function coupling can provide context to understand novel electrophysiological findings, such as iEEG measures of theta frequencies by age, based on well-documented age-related variability in regional brain structure ^38–40^. Based on initial evidence of age-related variability in peak theta frequencies ^9,30^ and well-documented evidence that the hippocampus matures earlier than association regions ^39,41–44^, such as the PFC, we hypothesized: (a) in association cortices, fast theta oscillations speed up and slow theta oscillations slow down with age into young adulthood; (b) in the hippocampus, fast theta frequencies predict memory independent of age; (c) attentional state (task-based vs. task-free) modulates age effects observed in (a) and (b), and; (d) age-related differences in theta frequencies vary by regional GMV.

**Figure 1.**
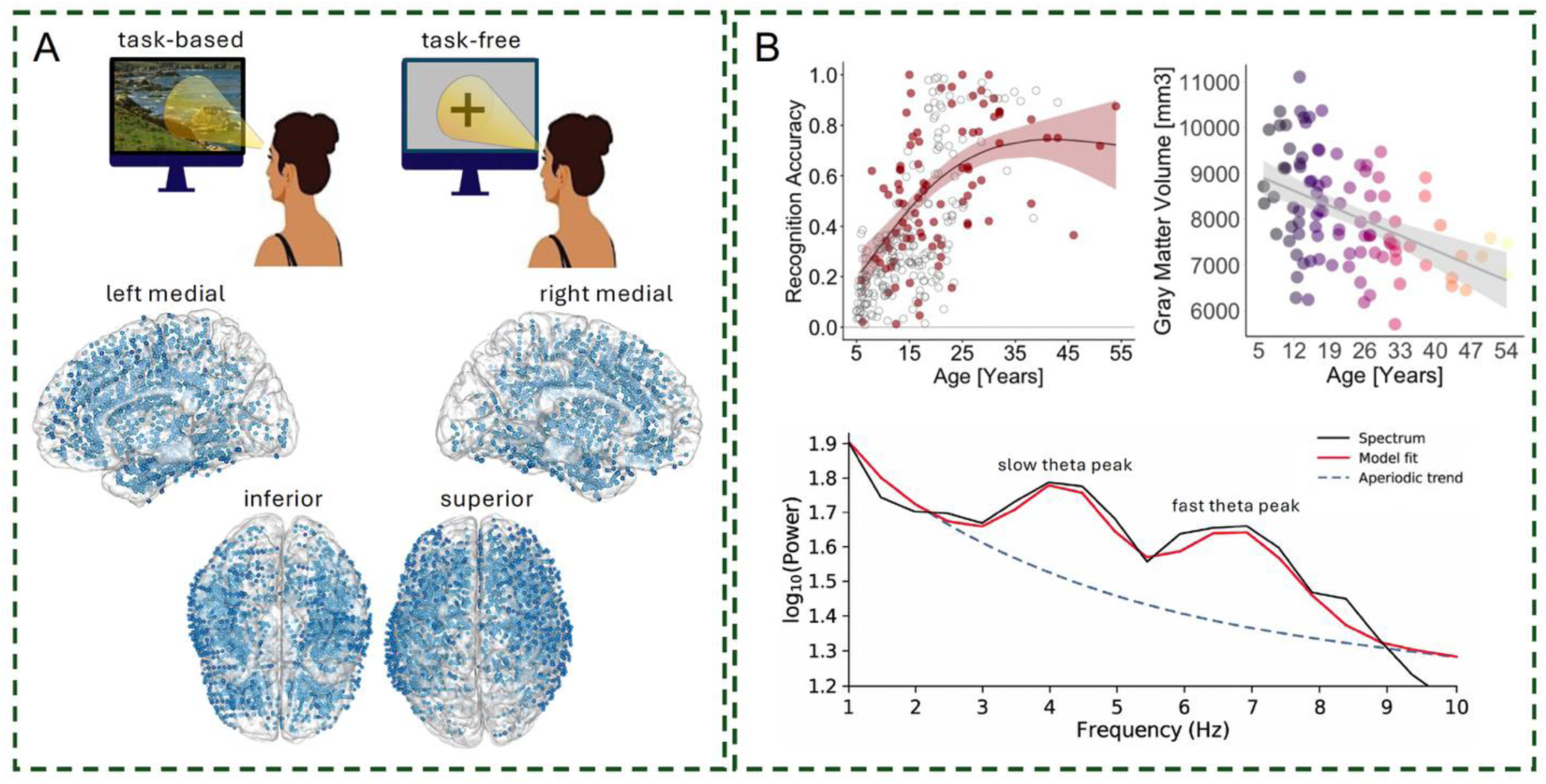
Design, channel coverage, and key variables. **(A)** Top: Intracranial neurophysiological activity was recorded during both task-based (top left) and task-free wake states (top right). Bottom: Seizure- and artifact-free intracranial channel placements (*n* = 5691) across all patients (*n* = 101) in MNI space. **(B)** Schematic of key dependent and independent variables. Top left: iEEG patients (red; *n* = 81) show the expected developmental trajectory of improved memory recognition from ∼5 – 30 years of age (one-sided non-linear regression, *p <*0.001) and fall in the range of age-matched, healthy controls (gray; *n* = 221). Shading indicates 83% CIs. Top right: age-related differences in global GMV (mm^3^) in our cohort, showing the expected developmental trajectory of decreased GMV from ∼5 – 54 years of age (one-sided linear regression, *p* <0.001). Bottom: exemplar power spectral density plot illustrating peak slow and fast theta frequencies over and above the aperiodic (1/ƒ-like) component

We first revealed dissociable slow and fast theta oscillations across the brain and uncovered a spatial gradient whereby slow theta is slowest in posterior regions and fastest in anterior regions, while the opposite was observed for fast theta. We then established developmental trajectories of slow and fast theta frequencies. Within regions, we found maximal separation between slow and fast theta sub-bands around 18-20 years of age. Between regions, interregional differences in slow theta also emerged around 20 years of age. By contrast, interregional differences in fast theta remained relatively stable across age. We further revealed how attentional state modulated age effects in select regions, including the hippocampus and PFC, and established predictive links between task-based and task-free slow and fast theta frequencies and individual memory outcomes. Notably, although hippocampal slow theta frequencies varied with age, slow theta frequencies were positively associated with memory performance after accounting for age, not fast theta oscillations as hypothesized. Finally, we observed no significant associations between age, frequency, and GMV, contrary to our hypothesis that age-related differences in theta frequencies would vary by regional GMV. Collectively, our findings establish developmental trajectories of slow and fast theta oscillations in localized brain regions, offer critical insights into the complex interplay of theta and behavior, and reveal how the maturation of slow theta oscillations in the hippocampus and PFC supports the development of attention and memory.

## Results

### iEEG memory and brain volume measures generalize to healthy populations

One hundred and one neurosurgical patients participated (mean age = 19.25, range = 5.93 – 54.00 years; 63 males). Patients were selected based on above-chance behavioral performance on two visual memory recognition tasks (mean normalized accuracy = 0.54, SD = 0.25, range = 0.01 – 1.00; *β* = 0.54, SE = 0.02, *p* <0.001) and/or if there was a task-free recording available. Those with major lesions, prior surgical resections, noted developmental delays, or neuropsychological memory test scores <80 were considered ineligible. We recently demonstrated ^33^ using a nonlinear regression (single spline with two internal knots) that there is a positive association between recognition accuracy and age (first knot: *β* = 0.90, SE = 0.16, *p* <0.001; second knot: *β* = 0.30, SE = 0.13, *p* = 0.027; *R^2^* = .27; see Figure 1C), indicating that iEEG patients exhibit the expected developmental trajectory of improved memory from age 5-30 years, consistent with age-matched, healthy controls ^3,9,45^. Analysis of global GMV by age indicated a negative association (*β* = -135.14, SE = 52.74, *p* =0.01, *R^2^* = .25; Figure 1C), indicating that with every one-year increase in age there is a 135mm^3^ reduction in GMV ^33^ after controlling for total intracranial volume. This further demonstrates that iEEG patients show the expected developmental trajectory of decreased GMV from age 5 to 54 years, consistent with well-documented decreases in GMV from childhood through adulthood in healthy individuals ^38–40,46^. These demonstrations support the idea that our iEEG analyses generalize to healthy populations ^47,48^.

### Peak theta frequencies differ by brain region

Before testing hypotheses, we first characterized regional differences in slow and fast theta frequencies by implementing linear mixed-effects models, regressing region onto slow and fast peak theta frequencies while regressing out attentional state (task-based, task-free) and age, treating participants and nested channels as random intercepts ^48^. Regions of interest (ROIs) were defined based on the Desikan-Killiany-Tourville (DKT) atlas ^49^ (for a summary of theta frequencies by brain region, see S1 – S4 in the supplementary material). We revealed a posterior-to-anterior gradient for slow theta, with slow theta oscillations slowest in posterior and fastest in anterior regions (χ2(20) = 67.04, *p* ≤ 0.001; Figure 2A). This pattern was opposite for fast theta, whereby fast theta oscillations were slowest in anterior and fastest in posterior regions (χ2(20) = 231.54, *p* ≤ 0.001; Figure 2B). Our data provide the first demonstration of dissociable slow and fast theta sub-bands across the whole brain.

**Figure 2.**
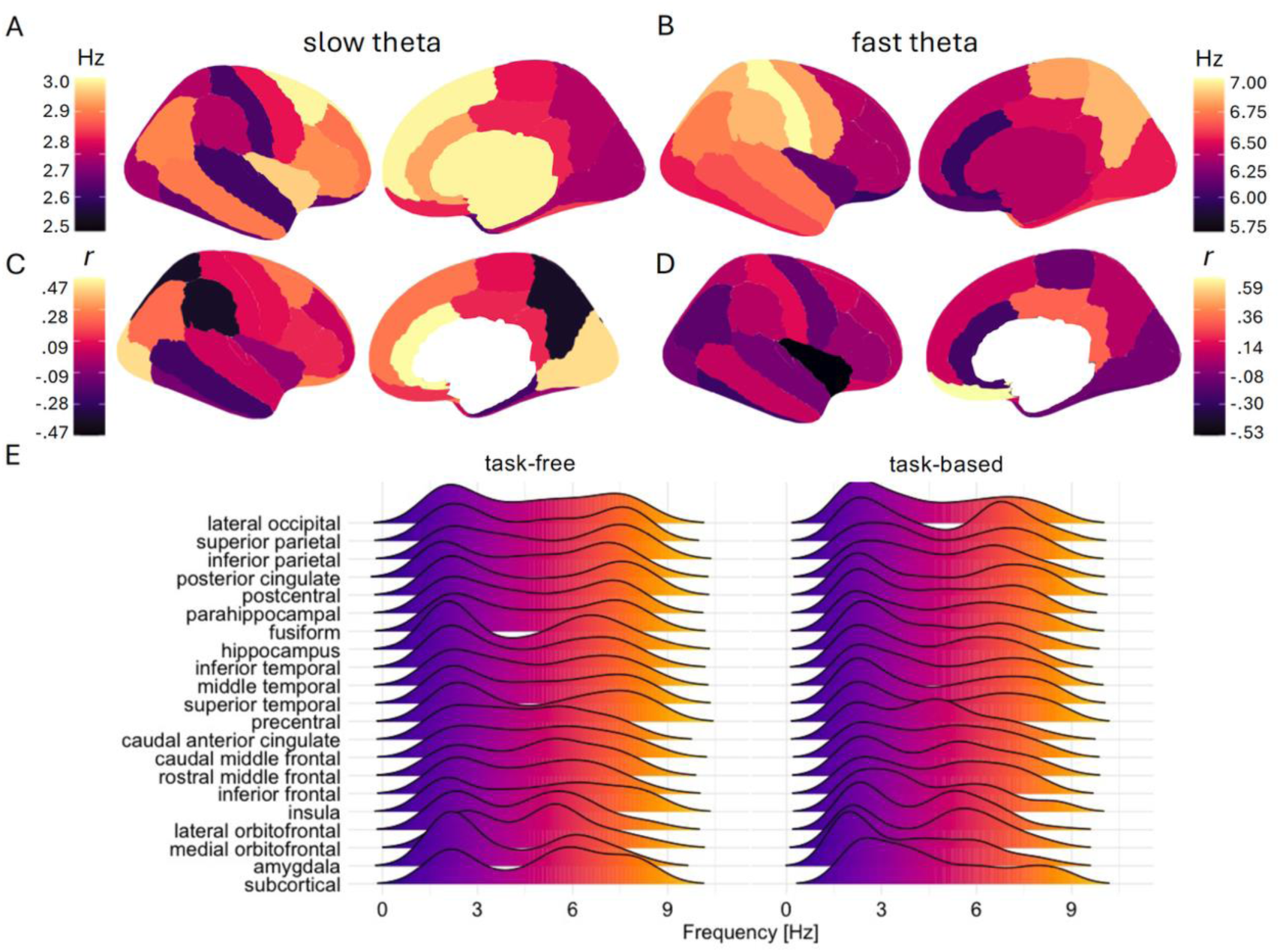
Slow and fast peak theta frequencies differ across the brain. Top row: Brain-wide standardized means (predicted marginal means) of regional slow (**A**; left) and fast (**B**; right) peak theta frequencies. Warmer colors/higher values indicate faster frequencies. Middle row: brain-wide correlations (Pearson *r*) between regional GMV (mm^3^) and slow (**C**; left) and fast (**D**; right) peak theta frequencies. Warmer colors/higher values indicate positive correlations and cooler colors/lower values indicate negative correlations. Note that the area corresponding to subcortical space is white as no analysis of subcortical GMV was performed. (**E**) Ridgeline plot illustrating the distribution of slow and fast peak theta frequencies (x-axis; higher values denote a faster frequency) by region (y-axis) and condition (left: task-free; right: task-based) according to an approximate posterior-to-anterior gradient.

Second, to characterize relationships between regional GMV and peak fast and slow theta frequencies (i.e., structure-function coupling), we implemented linear mixed-effects models, regressing region and GMV onto fast and slow theta frequencies, while regressing out attentional state and age, treating participants and nested channels as random intercepts. Controlling for age and attentional state, there were no significant relationships between fast or slow theta frequencies and regional GMV, despite large effect sizes (*r* range = -.53 - .59; Figure 2C and 2D).

### Slow and fast theta frequencies are most separated in young adulthood

Next, based on initial evidence that the theta band separates into slow and fast sub-bands in PFC ^9^, we examined hypothesis (a), that in association cortices, fast theta oscillations speed up and slow theta oscillations slow down with age into young adulthood. We further sought to establish the age in which slow and fast theta frequencies are most separated. We implemented a non-linear mixed-effects regression, modelling frequency as a function of band (slow and fast theta), age (fit with one spline; two knots), and region type (association, MTL, sensorimotor; see Table S5 for a summary of association, MTL, and sensorimotor regions), treating subject and DKT region as random effects on the intercept, and channel nested under subject. Although the age × region type × band interaction was non-significant (χ2(4) = 5.06, *p* = 0.28), we revealed a significant age × band interaction (χ2(2) = 12.04, *p* = 0.002; Figure 3A), whereby slow theta frequencies slow gradually beginning in childhood, and fast theta frequencies are stable across childhood and then slow with advancing age. Peak separation between slow and fast theta frequencies was observed around 18-20 years of age (Figure 3B). In addition, a significant region type × band interaction (χ2(2) = 36.15, *p* ≤ 0.001) revealed differences in the separation between slow and fast theta frequencies across region types (Figure 3C). This separation (fast − slow) was greatest in sensorimotor cortices relative to both association cortices and the MTL. At the whole-brain level, slow and fast theta frequencies showed increasing divergence across development into young adulthood, providing partial support for hypothesis (a). Regionally, however, this separation was not uniform, indicating maximal separation between theta sub-bands in sensorimotor cortices irrespective of age.

**Figure 3.**
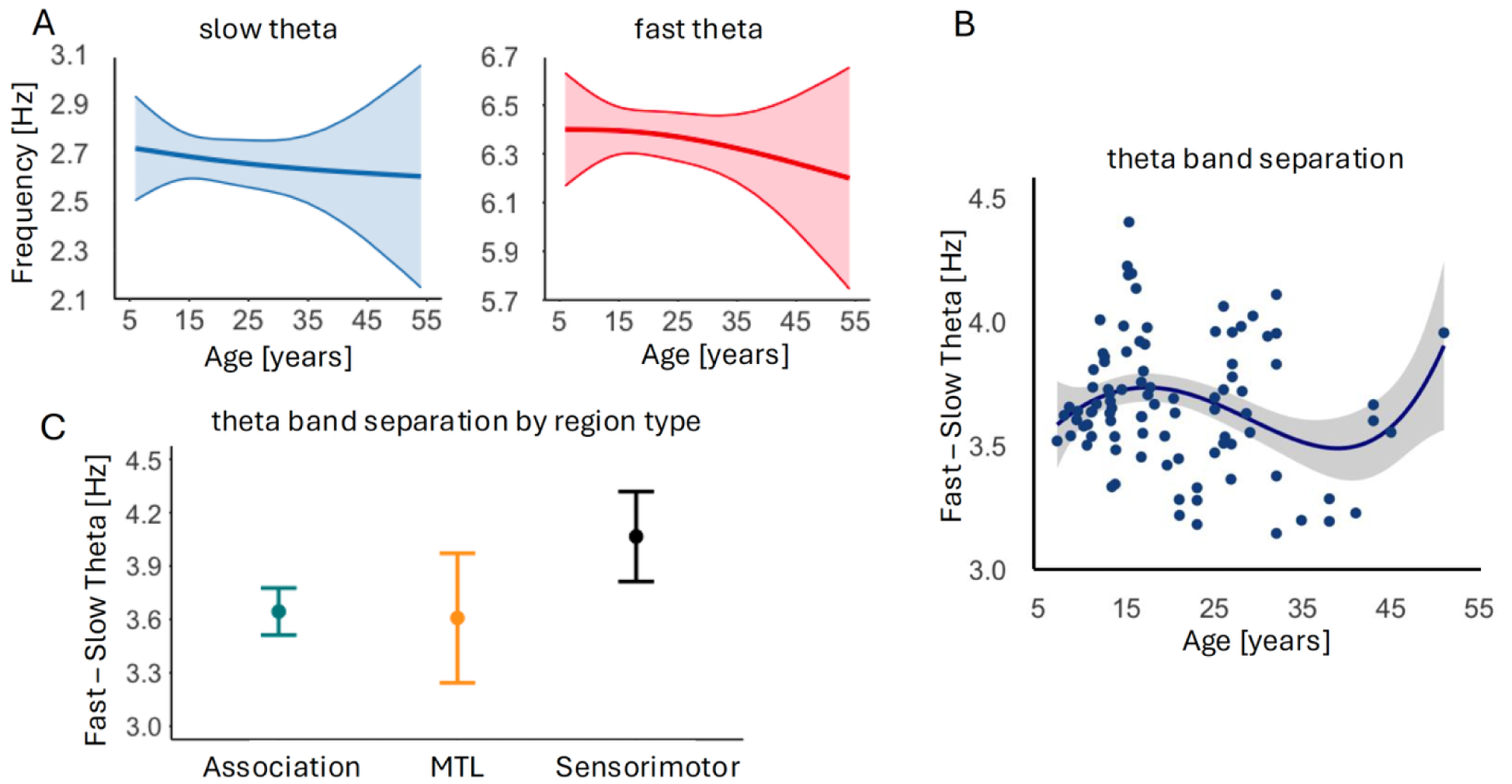
Slow and fast theta oscillations separate from childhood into adolescence. **(A)** Modelled effects for differences in slow and fast theta frequencies (y-axis; higher values denote a faster theta frequency) and age (x-axis). Shading indicates 83% CIs. **(B)** Separation between fast and slow theta frequencies (fast – slow; y-axis) as a function of age (x-axis). Data points represent moving averages of differences between fast and slow peak theta frequencies. The solid line represents a smoothed trend (locally estimated scatterplot smoothing; LOESS), reflecting the overall pattern in the data rather than subject-level differences. The seeming increase in frequency separation in older ages should be interpreted with caution because it is a likely artifact of fewer observations in this range. **(C)** Separation between fast and slow theta frequencies (fast – slow; y-axis) by region type. Whiskers indicate 83% CIs.

To further examine hypothesis (a), we implemented nonlinear mixed-effects regressions by sub-band, modelling slow and fast theta frequencies as a function of age (fit with one spline; two knots) and region type (association, MTL, sensorimotor), treating subject and DKT region as random effects on the intercept, and channel nested under subject. For slow theta, we revealed a significant age × region type interaction (*β* = 0.75 [95% CI = 0.31, 0.82], SE = 0.36, *p* = 0.037), revealing that slow theta begins to speed up in the MTL around 20 years of age, diverging from the gradual slowing of slow theta oscillations in association and sensorimotor cortices (Figure 4A). For fast theta, although there was neither an interaction of age and region type (*β* = 0.50 [95% CI = -0.46, 1.46], SE = 0.49, *p* = 0.31) nor a main effect of age (*β* = -0.23 [95% CI = -1.16, 0.67], SE = 0.46, *p* = 0.61), we confirmed a significant main effect of region type, whereby fast theta frequencies were fastest in sensorimotor cortices and slowest in the MTL (*β* = -0.77 [95% CI = -0.65, 0.64], SE = 0.26, *p* = 0.01; Figure 4B). These results indicate that rather than both sub-bands differing in opposite directions, slow and fast theta frequencies follow dissociable developmental trajectories, with slow theta exhibiting age-dependent modulation by region and fast theta exhibiting relatively age invariant regional effects (for model diagnostics, see Figures S6 and S7 in the supplementary material for slow and fast theta models, respectively).

**Figure 4.**
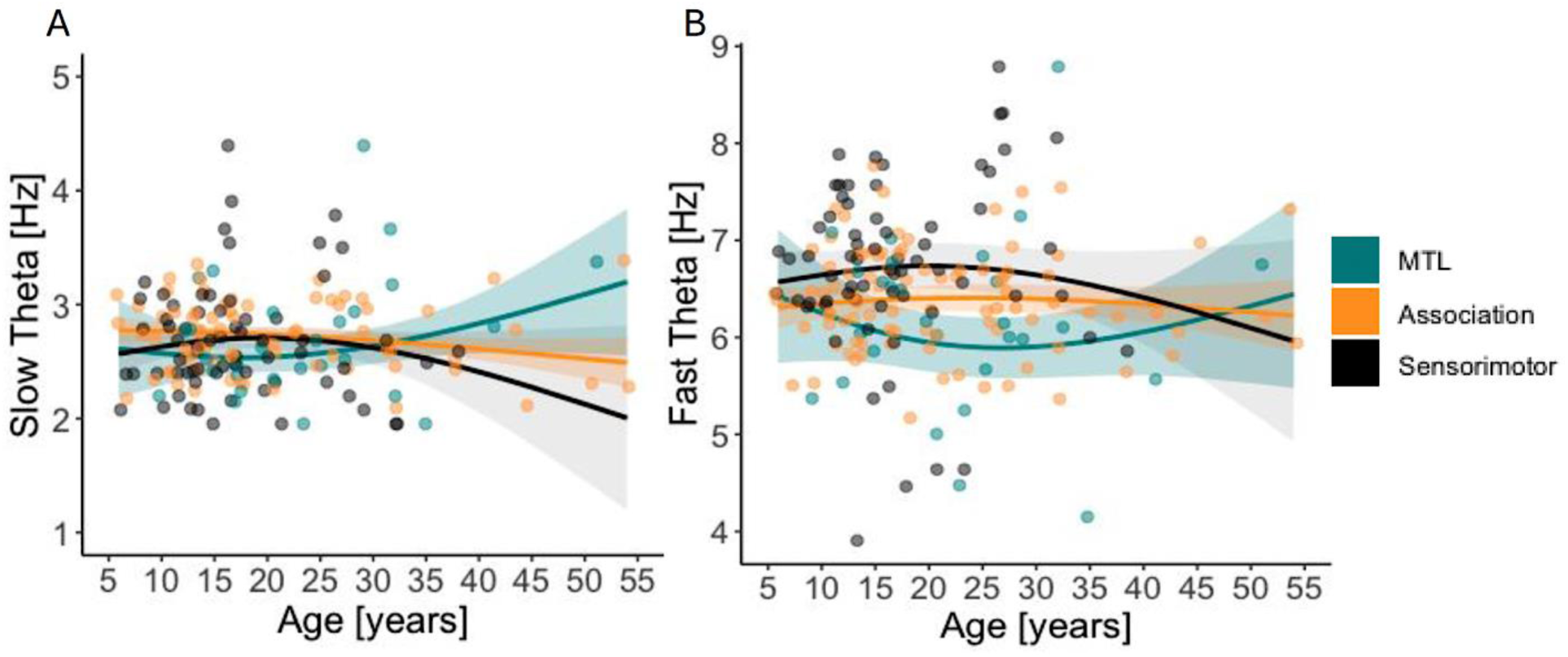
Aged-related differences in theta frequencies differ between association and sensorimotor cortices and the MTL. **(A)** Modelled effects for differences in slow theta frequencies (y-axis; higher values denote a faster slow theta frequency) and age (x-axis). **(B)** Modelled effects for differences in fast theta frequencies (y-axis: higher values denote a faster fast theta frequency) and age (x-axis). In both **(A)** and **(B)**, MTL regions are presented in teal, association cortices in orange, and sensorimotor cortices in black. Shading indicates 83% CIs. Individual data points represent peak slow and fast theta frequency values per participant averaged over channels.

### Regional theta frequencies differ by age and attentional state

Having demonstrated that slow and fast theta frequencies differentially relate to age across association, MTL, and sensorimotor regions, we next sought to establish developmental trajectories of theta frequencies within localized brain regions and test hypothesis (c), that age effects would differ between attentional states. We follow by testing hypothesis (b) through systematic investigation of associations between age, task-based and task-free theta frequencies, and memory outcomes. To identify regional age effects in slow and fast theta frequencies and whether they differ by attentional state, we implemented separate linear mixed-effects models for each ROI. Our strategy for each analysis was to fit a model to the dependent variable of interest (i.e., slow or fast peak theta frequencies) and regress the estimates onto age, attentional state (task-based, task-free), and the interaction of age and attentional state. All models were fit with by-participant and by-task random intercepts, with channel nested under participant.

For slow theta, we revealed significant age × attentional state interactions in caudal middle frontal gyrus (cMFG; *β* = -0.01 [95% CI = -0.02, -0.003], SE = 0.005, *p* = 0.009) and the hippocampus (*β* = 0.03 [95% CI = 0.004, 0.06], SE = 0.01, *p* = 0.02; Figure 5A, 5B). In cMFG, task-free oscillations were faster than task-based oscillations in children, and the opposite was observed in adults; the pattern reversed around 15 years of age (Figure 5B left). In the hippocampus, by contrast, task-based oscillations were faster than task-free oscillations in children and the opposite was observed in adults (Figure 5B right); the pattern reversed around 20 years of age. For visualization of all main effects and model diagnostics, see Figures S8 – S11 in the supplementary material.

**Figure 5.**
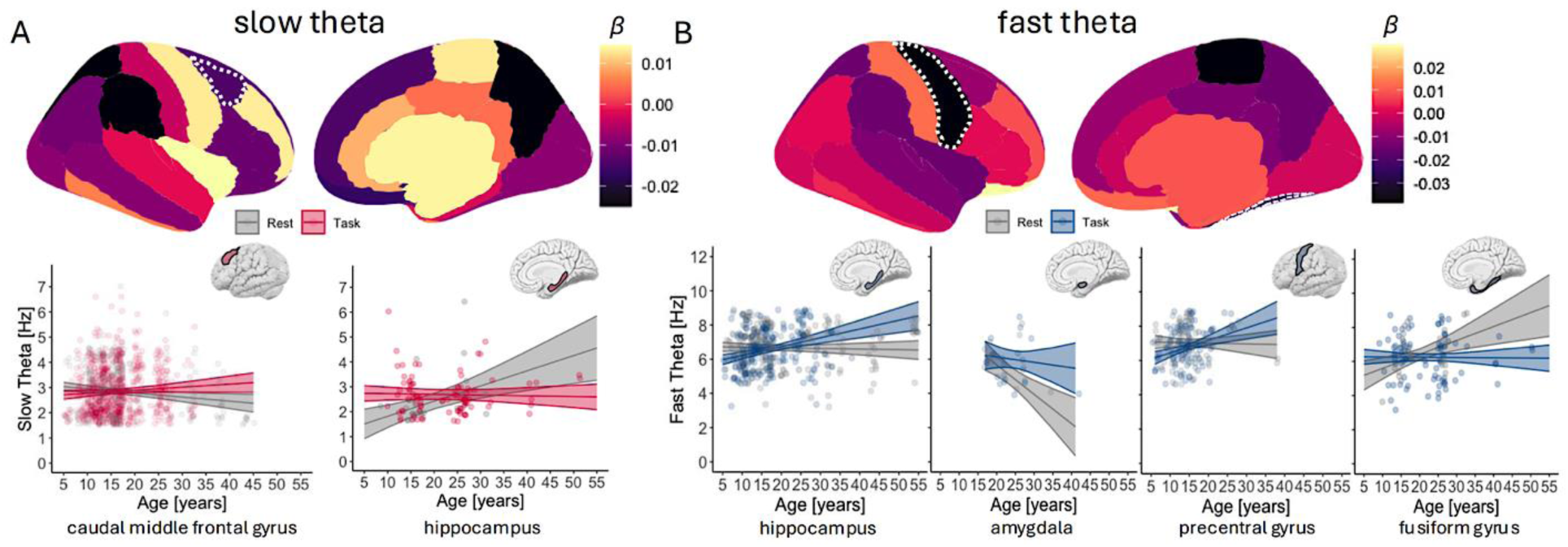
Slow and fast theta frequencies differ by age and attentional state. **(A)** Top row: Brain-wide age and condition interactions on regional peak slow theta frequencies. Regions with statistically significant interactions between age and attentional state (*FDR* < 0.05) are indicated by dashed borders. Bottom row: scatterplots illustrating interactions between age (x-axis; in years) and attentional state (red = task-based; gray = task-free) on peak slow theta frequencies (y-axis; higher values denote a faster slow theta frequency) in regions with statistically significant interactions. Individual data points represent single participant data averaged across channels for each representative ROI. Shading indicates 83% CIs. **(B)** Same as (**A**) for peak fast theta frequencies.

For fast theta, we observed significant age × attentional state interactions in the hippocampus (*β* = 0.04 [95% CI = 0.003, 0.08], SE = 0.02, *p* = 0.03), amygdala (*β* = -0.07 [95% CI = -0.12, -0.02], SE = 0.02, *p* = 0.01), precentral gyrus (*β* = -0.03 [95% CI = -0.07, -0.002], SE = 0.02, *p* = 0.03), and fusiform gyrus (*β* = -0.03 [95% CI = -0.04, -0.01], SE = 0.008, *p* <0.001; Figure 5C, 5D). Differential patterns were observed across temporal ROIs, with task-based oscillations faster than task-free oscillations in the adult hippocampus and amygdala, and task-free oscillations faster than task-based oscillations in the adult fusiform gyrus. Differences between task-states emerged around 20 years of age across temporal ROIs. In the precentral gyrus, task-free oscillations were faster than task-based oscillations in children and the opposite was observed in adults; the pattern reversed around 20 years of age. Taken together, these results demonstrate that age effects in slow and fast theta frequencies are modulated by attentional state in a region-specific manner, supporting our hypothesis. For visualization of all main effects and model diagnostics, see Figures S12 – S14 in the supplementary material.

The hippocampus is the only region that showed developmental sensitivity to attentional state across sub-bands. For slow theta, hippocampal peak frequencies increased with age preferentially during task-free rest, with the direction of task-free versus task-based differences reversing in young adulthood. A different pattern was observed for fast theta, where hippocampal peak frequencies increased with age preferentially during task engagement, with the direction of differences again reversing in young adulthood. Together, these findings indicate that the trajectory of hippocampal theta maturation differs between slow and fast theta sub-bands according to attentional state, revealing functional theta maturation into young adulthood despite stable hippocampal volume across this age range ^44^. In parallel, the PFC (here, cMFG) showed age-dependent reversals in slow theta as a function of attentional state, suggesting that the developmental reorganization of frontotemporal theta dynamics contributes to age-related differences in attention.

### Task-free and task-based slow and fast theta frequencies differentially predict memory

Having demonstrated that memory performance improves with age, with marked variability among adolescents (Figure 1C), we examined whether age interacts with regionally specific task-based and task-free slow and fast theta frequencies, respectively, to predict memory performance. For each analysis, we fitted a general linear model to recognition accuracy and regressed the estimates onto age and slow or fast theta frequencies (task-based or task-free), and the interaction of age and frequency. In line with hypothesis (b), this analysis specifically evaluated whether hippocampal fast theta frequency predicted recognition accuracy independent of age.

For task-based slow theta (Figure 6A-B), we revealed a main effect of frequency in the hippocampus (*β* = 1.00 [95% CI = 0.25, 2.16], SE = 0.42, *p* = 0.02; *R*^2^ = .34), such that memory performance increased with faster slow theta oscillations independent of age (Figure 6B left), linking hippocampal slow theta oscillations to memory as well as attentional state. There was also a significant frequency × age interaction in the inferior frontal gyrus (IFG; *β* = -0.05 [95% CI = -0.09, 0.005], SE = 0.02, *p* = 0.02; *R*^2^ = .35; Figure 6B right), such that slower slow theta oscillations were associated with worse memory outcomes in children but better memory in adults. For task-free slow theta (Figure 6C-D), we observed a significant frequency × age interaction in the fusiform gyrus (*β* = -0.05 [95% CI = 0.004, 0.09], SE = 0.02, *p* = 0.01; *R*^2^ = .22), such that slower slow theta oscillations were associated with better memory outcomes in children but worse memory in adults. For model diagnostics for task-based and task-free slow theta on memory, see Figures S15 – S20 in the supplementary material.

**Figure 6.**
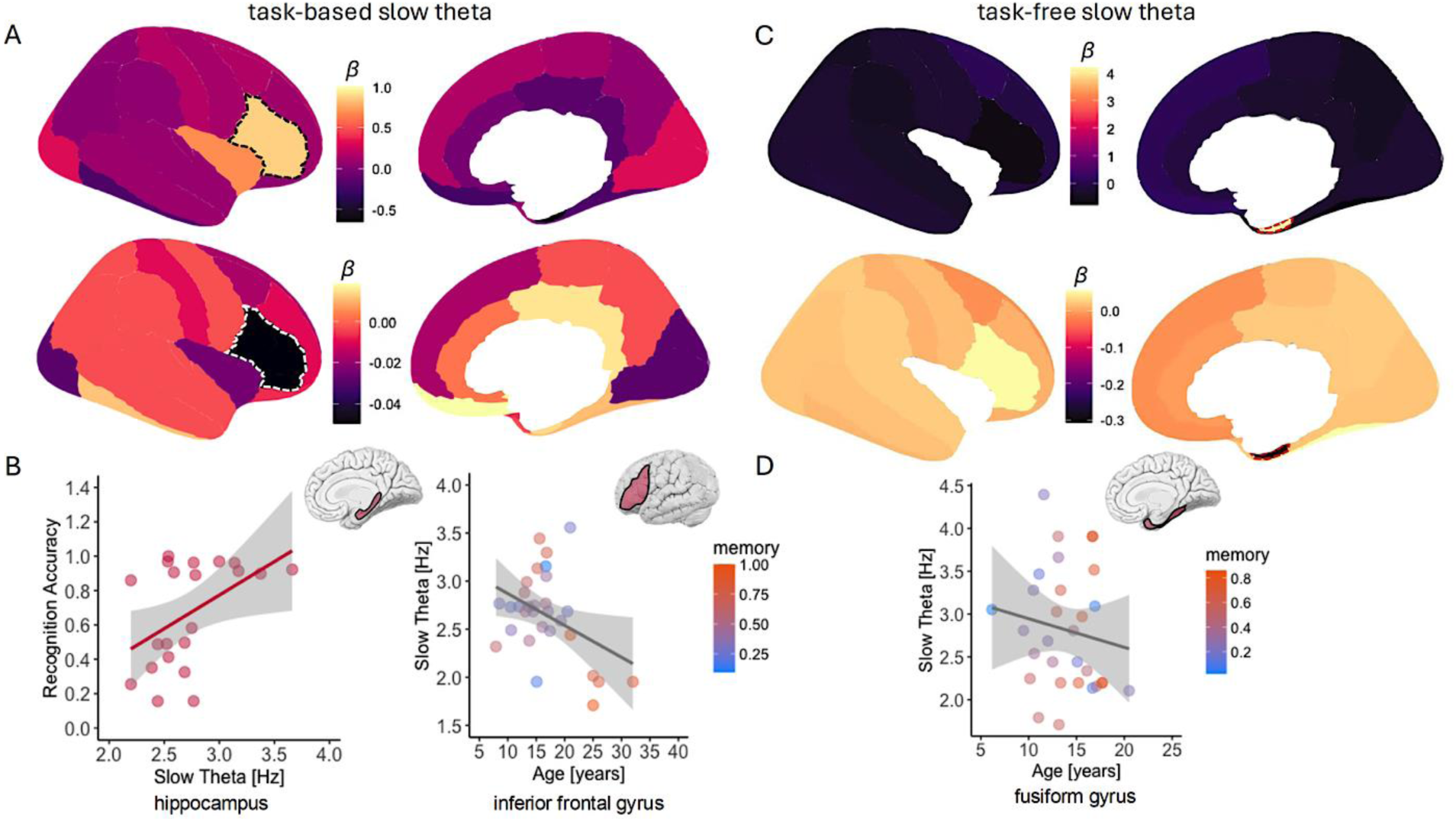
Task-based and task-free regional slow theta frequencies predict memory performance. **(A)** Top, brain-wide maps demonstrating the main effect of frequency, and bottom, age × frequency interaction on memory for task-based slow theta frequencies. The color bars denote the unstandardized beta coefficients for the main and interaction effects. Regions which had a significant main effect or interaction (i.e., *p* <.05) are presented in dashed borders. Regions in white did not have sufficient data points (e.g., insufficient detectable peak oscillations) and thus models did not converge. **(B)** Left: scatterplot displaying the significant main effect of frequency independent of age on memory performance in the hippocampus for task-based slow theta frequencies. Right: scatterplot displaying the significant age × frequency interaction on memory performance in the inferior frontal gyrus for task-based slow theta frequencies. Shading indicates 83% CIs. **(C)** Same as (**A**) for task-free (resting-state) slow theta frequencies. **(D)** Scatterplot displaying the significant age × frequency interaction on memory performance in the fusiform gyrus for task-free slow theta frequencies, same conventions as (**B**).

For task-based fast theta (Figure 7A-B), we revealed main effects of frequency in the lateral orbitofrontal cortex (lOFC; *β* = 0.77 [95% CI = 0.03, 1.73], SE = 0.37, *p* = 0.04; *R*^2^ = .30) and middle temporal cortex (*β* = 0.17 [95% CI = -0.01, 0.008], SE = 0.08, *p* = 0.04; *R*^2^ = .42). In both regions, memory increased with faster oscillations independent of age. For task-free fast theta (Figure 7C-D), we observed significant frequency × age interactions in the anterior cingulate cortex (ACC; *β* = 0.08 [95% CI =-0.07, 0.16], SE = 0.02, *p* = 0.01; *R*^2^ = .48) and middle temporal cortex (*β* = 0.03 [95% CI = -0.01, 0.05], SE = 0.01, *p* = 0.006; *R*^2^ = .40). In both regions, children who had slower task-free fast theta oscillations demonstrated superior memory performance, while adults with slower task-free fast theta oscillations demonstrated inferior memory performance. These results demonstrate that both task-based and task-free slow and fast theta frequencies, in some cases independent of age, predict memory outcomes in a region-specific manner. For model diagnostics for task-based and task-free fast theta on memory, see Figures S21 – S26 in the supplementary material.

**Figure 7.**
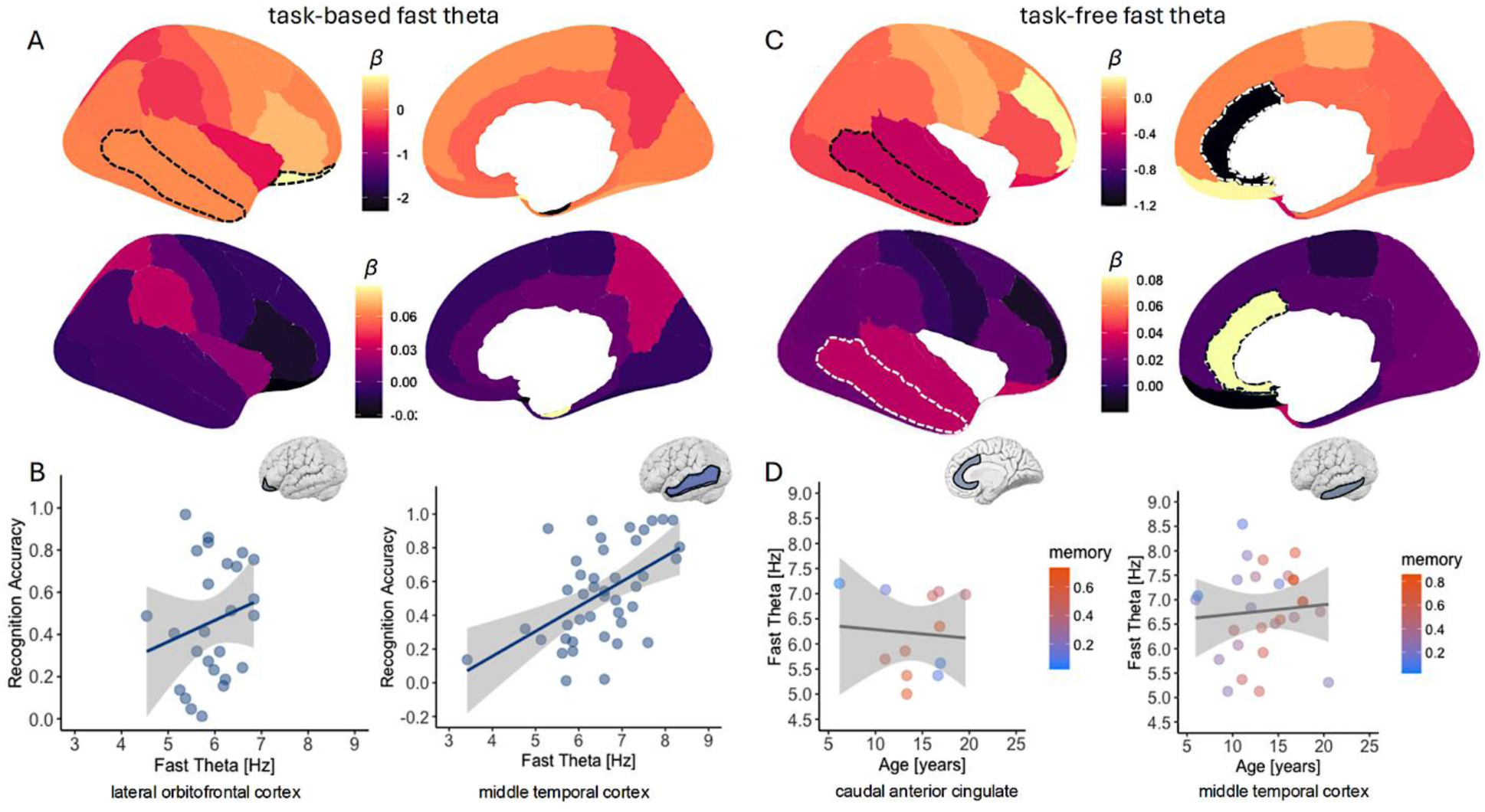
Age-related task-based and task-free regional fast theta frequencies predict memory performance. **(A)** Top, brain-wide maps demonstrating the main effect of frequency and bottom, age × frequency interaction on memory for task-based fast theta frequencies. The color bars denote the unstandardized beta coefficients for the main and interaction effects. Regions which had a significant main effect or interaction (i.e., *p* <.05) are presented in dashed borders. Regions in white did not have sufficient data points (e.g., insufficient detectable peak oscillations) and thus models did not converge. **(B)** Scatterplots displaying the significant main effect of frequency independent of age on memory performance in lateral orbitofrontal cortex (left) and middle temporal cortex (right) for task-based fast theta frequencies. Shading indicates 83% CIs. **(C)** Same as (**A**) for task-free (resting-state) fast theta frequencies. **(D)** Scatterplots displaying the significant age × frequency interactions on memory performance in caudal anterior cingulate and inferior temporal cortex for task-free fast theta frequencies, using the same conventions as (**B**).

Together, these findings indicate a dissociation between hippocampal and prefrontal theta dynamics during task engagement across development. Contrary to hypothesis (b), that age-invariant faster hippocampal fast theta frequencies would be associated with superior memory, we revealed that hippocampal age-invariant faster *slow* theta frequencies during task engagement were associated with superior memory. We further highlight age-dependent effects in the PFC during task engagement (here, IFG), where the age-related slowing of slow theta oscillations was associated with age-related improvements in memory from childhood into adulthood. Additional effects were observed in other frontotemporal regions during task engagement and task-free rest, demonstrating direct relationships between theta frequencies and memory outcomes across the frontotemporal network known to support memory function. Frontotemporal theta oscillations may thus serve as key markers of memory development.

### Regional gray matter volumes do not explain age-related differences in theta frequencies

Thus far, we have established that frontotemporal slow and fast theta frequencies differ by age and attentional state, and predict age-related variability in memory outcomes. Last, we focus on structure-function relationships by testing hypothesis (d), that age-related differences in slow and fast theta frequencies vary by regional GMV. Importantly, we recently demonstrated in this cohort that there are significant age-related reductions in global GMV (see Figure 1C) and in GMV across temporal, frontal, and parietal ROIs ^33^, replicating previous reports of age-related reductions in GMV in the broader population ^38–40^. Notably, we did not observe an age effect on hippocampal GMV, further replicating previous reports of stable hippocampal volume from age five into adulthood ^44^.

We fit mixed-effects models to task-based and task-free slow and fast theta frequencies and regressed these estimates onto age, GMV, and the interaction between age and GMV, while controlling for total intracranial volume. All models were fit with by-participant and by-task random intercepts, with channel nested under participant. We observed no significant effects for either task-based or task-free slow or fast theta frequencies, suggesting that the development of theta frequencies is independent of regional GMV.

## Discussion

We mapped slow (∼1.5–4.5 Hz) and fast (∼4.5–9 Hz) theta oscillations across the human brain from childhood to late middle adulthood. Our results demonstrated: (I) a posterior-to-anterior gradient in theta oscillations, with slow theta slowing down and fast theta speeding up across the cortical axis, establishing distinct regional profiles for slow and fast theta sub-bands (Figure 2); (II) maximal separation between slow and fast theta frequencies around 18-20 years, suggesting a developmental specialization of slow and fast theta bands (Figure 3); (III) regional and state-dependent age effects on theta frequencies, with attentional state modulating age effects on slow theta in the hippocampus and PFC (Figure 5); (IV) regional and state-dependent frequency effects on memory performance, with task-based slow theta predicting memory independent of age in the hippocampus and as a function of age in the PFC (Figure 6); and (V) no significant associations between theta frequencies, GMV, and age. Collectively, these findings link slow theta oscillations in the hippocampus and the PFC to the maturation of attention and memory and establish frontotemporal theta oscillations as key markers of memory development, independent of regional brain structure (for a summary of key results, see Figure 8).

**Figure 8.**
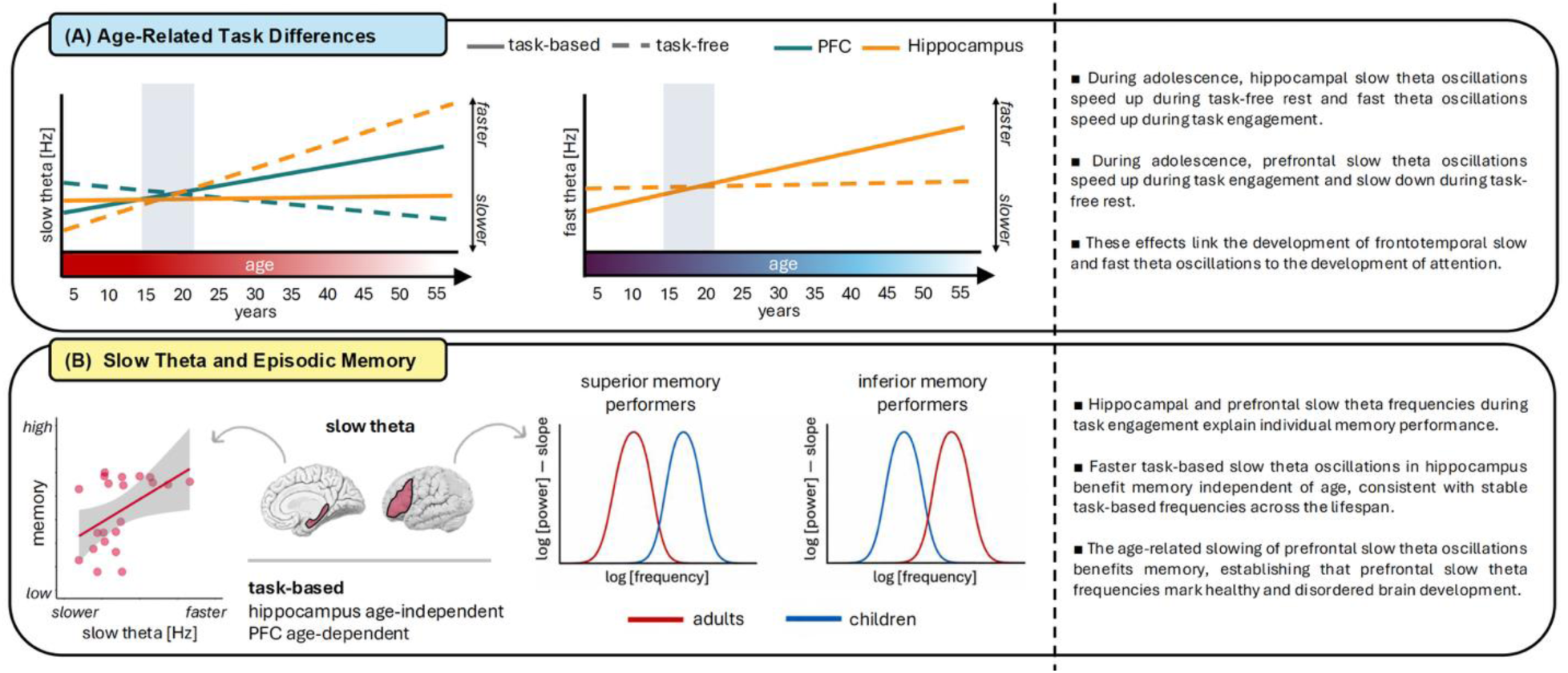
Slow and fast theta frequencies in the hippocampus and the PFC differ by age and attentional state, and slow theta frequencies predict individual memory. **(A)** Slow and fast theta frequencies show region- and task-dependent developmental trajectories in the hippocampus and the PFC, linking the development of frontotemporal slow and fast theta oscillations to the development of attention. **(B)** Task-based slow theta frequencies predict individual differences in memory, with age-independent effects in the hippocampus and age-dependent effects in the PFC, linking prefrontal slow theta oscillations to brain development.

### Attention modulates slow theta frequencies by age in hippocampus and PFC

To date, most work on the development of theta oscillations has relied on scalp-EEG ^30–32^ (cf. ^9,50^), reporting that a posterior theta oscillation speeds up across development and an anterior attentional control-related theta oscillation shows a more protracted developmental profile ^30^. Most work has not examined slow and fast theta oscillations separately, conflating their unique functions in cognitive performance (cf. ^2,51^). Here, by utilizing spatiotemporally precise iEEG recordings from task engagement and task-free rest, we reveal that attentional state modulated age-related differences in slow and fast theta frequencies in a regionally specific manner. In the hippocampus, slow theta was faster during task engagement in childhood, with this pattern reversing by adulthood due to an age-related speeding of task-free oscillations. In contrast, in the PFC (here, cMFG), slow theta was faster during task-free rest in childhood, with this pattern reversing by adulthood due to an age-related speeding of task-based oscillations. These opposing patterns indicate that hippocampal and prefrontal slow theta oscillations do not follow a common developmental trajectory. These findings establish that the influence of attentional state on slow theta oscillations differs across regions and ages, suggesting that functional maturation of the hippocampus and the PFC underpins age-related differences in the temporal organization of neural activity that supports attention and memory.

Notably, attentional effects consistently reversed around late adolescence to early adulthood across frontotemporal regions, coinciding with the age of maximal separation between slow and fast theta frequencies across the brain. This convergence is consistent with the well-established finding that adolescence is a period of functional specialization, during which neural systems become increasingly differentiated ^33,52,53^. Accordingly, our findings reveal that frontotemporal theta frequencies similarly differentiate by attentional state during adolescence. This developmental transition closely mirrors our recent observation that aperiodic activity in the PFC diverges according to attentional state during adolescence, with the direction of task-state differences reversing around 18-20 years of age, consistent with the development of cognitive control ^33^. Across both oscillatory and aperiodic neural activity, adolescence appears to mark a critical period in which prefrontal signals differentiate and stabilize, potentially supporting the transition to “adult-like” attentional and mnemonic abilities.

There was also a difference in fast theta as a function of age and task state across frontotemporal regions, including the hippocampus, with frequencies in the amygdala and fusiform gyrus exhibiting opposite age by task state interactions relative to the hippocampus. One interpretation is that these state-dependent differences reflect developmental differences in how theta provides temporal structure for the encoding of mnemonic information and its transformation into lasting memory representations ^21,26^. Theta oscillations are proposed to provide optimal windows for neuronal firing, such that spikes occur preferentially at specific phases of the theta cycle ^26,28,54^, dictating when information can be transmitted and integrated ^55,56^. From this perspective, age-related differences in fast theta frequencies may influence the duration and stability of these temporal windows, with hippocampal fast theta oscillations dictating how task-relevant information is encoded into memory^25^.

### Task-based slow theta frequencies in hippocampus and PFC predict memory

Do age-related differences in slow and fast theta frequencies predict age-related differences in memory? Prior work has reported mixed findings when relating theta activity to different aspects of cognition ^57–59^. Both theta power and frequency have been linked to successful memory encoding and retrieval, particularly in task-based contexts ^59^. However, these effects are not uniform. In some studies, increased power and faster frequencies have been associated with improved memory performance, whereas others have reported opposing effects depending on task demands and the time windows analyzed (for reviews, see ^21,59^). Similarly, studies in young adults have shown that theta dynamics differentially support performance on decision-making ^60,61^ and learning ^29,62^ tasks, suggesting that the functional role of theta varies with cognitive demands and behavioral contexts. However, previous work has focused on either task-based or task-free theta activity in isolation, often within restricted brain regions or using scalp-EEG. As a result, it has remained unclear how slow and fast theta frequencies in different task states and brain regions relate to memory across the lifespan.

Here, we revealed that frontotemporal theta frequencies predict individual memory performance, illuminating how the pattern of slow theta effects during attention to to-be-remembered visual information differs between the hippocampus and PFC. In the hippocampus, faster task-based slow theta oscillations predicted better memory independent of age. This finding is inconsistent with our hypothesis that memory-related effects would be specific to fast theta. This hypothesis was based on prior reports that subsequently remembered items are associated with faster theta frequencies than subsequently forgotten items, together with evidence that hippocampal-cortical synchrony during encoding falls within the fast theta range ^2,63^. Instead, our results point to hippocampal slow theta oscillations being more strongly involved than fast theta oscillations during memory formation, independent of age. By contrast, prefrontal slow theta oscillations predicted individual memory outcomes by age, such that slower slow theta oscillations in the IFG were associated with worse memory outcomes in children but better memory outcomes in adults. Together, these findings indicate that hippocampal and PFC slow theta frequencies support memory through distinct mechanisms: faster hippocampal slow theta benefits memory irrespective of age, whereas the age-related slowing of prefrontal slow theta benefits memory by age. This dissociation links hippocampal slow theta frequencies to individual differences in memory and establishes prefrontal slow theta frequencies as a marker of brain development.

Prior work has also suggested that developmental differences in MTL and PFC theta frequencies contribute indirectly to memory through the developmental refinement of MTL-PFC interactions that support successful memory formation ^2,9^. One possibility is that the developmental refinement of slow theta supports memory by strengthening both local neuronal organization within and coordinated interactions between the hippocampus and PFC. Indeed, successful memory encoding is associated with increased temporal alignment of neural activity in the slow theta range ^55^ and superior memory performance is associated with stronger slow, but not fast, theta phase-locking between regions ^51^. From this perspective, the age-independent relationship between hippocampal slow theta and memory may reflect an early-established mechanism of temporal neuronal organization, whereby hippocampal neurons are more likely to fire at consistent time points within the theta cycle ^28,54^ in both adults and children. Conversely, age-dependent effects in the PFC likely reflect developmental refinements in how this temporal structure is recruited during higher-order cognitive operations, including the selection, maintenance, and monitoring of task-relevant information during memory encoding ^5,64^. Future work combining measures of frequency and interregional connectivity will be necessary to determine how these processes interact during the development of higher-order cognition.

In addition, faster fast theta oscillations during task engagement in lateral orbitofrontal and middle temporal cortices predicted memory independent of age, whereas faster fast theta oscillations during task-free rest in anterior cingulate and middle temporal cortices predicted better memory in adults but worse memory in children. Together, these findings suggest that fast theta supports memory through distinct frontotemporal circuits whose functional contributions become increasingly differentiated with development according to attentional state.

### The development of slow and fast theta frequencies is independent of regional grey matter volume

Despite clear age-related reductions in global and regional GMV ^33^, we found no evidence that slow and fast theta frequencies are associated with GMV, regardless of whether age was included in statistical models. Prior work has demonstrated that hippocampal volume remains relatively stable from childhood through early adulthood ^44^. Consistent with this observation, we observed age-related differences in hippocampal theta frequencies despite finding no relationship between theta frequencies and regional GMV. Our results therefore support and extend previous work by suggesting that developmental differences in hippocampal neural dynamics occur independently of gross structural maturation. One possibility is that theta frequencies reflect properties of neural timing that are more closely related to synaptic or circuit-level dynamics than to gross structural measures. For example, theta frequencies may reflect or be implicated in the balance between excitatory and inhibitory synaptic activity ^64,65^, the time constants of local recurrent circuits ^66^, or the degree of phase locking across neuronal populations ^22^, none of which are directly captured by volumetric measures such as GMV. From this perspective, the absence of a relationship between theta frequencies and GMV suggests that any synaptic and circuit-level dynamics reflected by theta frequencies are not readily indexed by gross volumetric measures, although theta frequencies may still relate to other structural or microstructural properties not captured by GMV ^67,68^ – a possibility that should be explored in future work.

### Limitations and future directions

We identified regionally specific age-related differences in slow and fast theta frequencies that depended on attentional state and predicted memory. However, several limitations should be considered. First, the present study was cross-sectional, and thus we cannot determine whether the observed age-related differences reflect within-subject developmental changes. Longitudinal studies will be necessary to establish these trajectories ^3^. Second, although our cohort exhibited expected developmental patterns in memory and brain structure consistent with healthy populations ^48^, use of a neurosurgical sample may still limit generalizability. Third, sampling was uneven across regions, with some regions having relatively low channel counts ^33^, and some channels did not exhibit a detectable slow or fast theta peak, further reducing statistical power and limiting interpretation of null findings. We caution against overinterpreting non-significant effects in sparsely sampled regions, such as the amygdala and insula, where lower electrode coverage reduced sensitivity to detect effects ^9,33,48^. Fourth, our analyses focused on peak frequency and did not examine time-resolved oscillatory dynamics, such as phase resetting, bursting, or interregional coupling. These temporal dynamics, particularly slow theta phase resetting and interregional coupling, are closely linked to successful memory encoding and may vary with age ^2,3,9,16^. As such, peak frequency should be interpreted as one component of theta dynamics, and future work integrating frequency with time-resolved measures would provide a more complete account of how theta supports memory development. Finally, although theta frequency development is independent of GMV, other biological factors, such as myelination, synaptic efficiency, or neuromodulatory systems, may shape oscillatory development.

### Implications

These findings address a longstanding gap in developmental neuroscience, as research has historically focused on young adults and provided limited insight into how oscillatory dynamics vary across the lifespan. By characterizing slow and fast theta frequencies from childhood to late middle adulthood, the present study provides a detailed account of how theta oscillatory frequencies relate to age-related differences in memory across development. Our findings thus have important implications for models of neurocognitive development ^2,3,52,69,70^.

Rather than reflecting a uniform maturation of oscillatory activity, theta frequencies show regionally and functionally specific developmental patterns, with slow theta oscillations in the hippocampus and the PFC playing a central role in the development of attention and memory. Improvements in memory across childhood and adolescence appear to be supported by age differences in the timing of neural activity, particularly prefrontal slow theta oscillations that organize neuronal firing during encoding ^55^. This interpretation is consistent with prior work suggesting that theta oscillations provide optimal temporal windows for spiking during memory encoding ^54^, and the infrastructure for interregional communication between the hippocampus and PFC ^9,22,26,66^. Understanding how age-related differences in theta frequencies relate to memory is important for understanding cognitive function in everyday contexts, given well-established differences in both brain structure and behavior across the lifespan. From this perspective, identifying typical patterns of theta development is also necessary to provide a benchmark for detecting atypical oscillatory dynamics. Clinically, our results may inform the early identification of neurodevelopmental and age-related conditions characterized by memory and attentional difficulties, including attention-deficit/hyperactivity disorder ^71,72^, developmental language disorders ^73^, and age-related cognitive decline ^74,75^.

### Conclusions

We revealed that slow and fast theta oscillations follow distinct developmental trajectories across the human brain, illuminating regional frontotemporal theta frequencies in the development of attention and memory. We further demonstrated how attentional state modulates age-related differences in slow theta oscillations in the hippocampus and the PFC, with opposing patterns across regions that reverse in adolescence, contributing to a growing body of literature on the development of “adult-like” cognitive abilities ^2,3,13,30,31,33,45,50,53^. We also established the functional relevance of slow theta during memory formation, illuminating prefrontal slow theta oscillations as a marker of brain development. Finally, we showed that developmental differences in theta frequency sub-bands are not explained by macroscopic brain structure. Taken together, these findings establish brain-wide patterns in theta frequencies, their relationship to individual differences in memory, and dissociable contributions of slow theta oscillations in the hippocampus and the PFC to the development of attention and memory across the lifespan.

## Methods

### Participants

Participants were 101 neurosurgical patients aged 5.93 – 54.00 years (63 males; mean age = 19.25) undergoing iEEG monitoring as part of clinical seizure management (see S27 in the supplementary material for a summary of participant characteristics). Patients with major lesions, prior surgical resections, noted developmental delays, or neuropsychological memory test scores <80 were considered ineligible. Patients were recruited from Northwestern Memorial Hospital, the Ann & Robert H. Lurie Children’s Hospital of Chicago, the Children’s Hospital of Michigan, the University of California (UC), San Diego Rady Children’s Hospital, UC Irvine Medical Center, UC Davis Medical Center, UC San Francisco Medical Center, Mount Sinai Hospital, California Pacific Medical Center, St. Louis Children’s Hospital, and Nationwide Children’s Hospital. The institutional review boards of Northwestern University (no. STU00215843), Lurie Children’s Hospital (no. 2022-5020), Wayne State University (no. 048404MP2E), UC Irvine and UC San Diego (no. 2014-1522), UC Davis (no. 1623773-1), UC San Francisco (no. 10-03842), Mount Sinai (no. STUDY-22-00529), California Pacific Medical Center (no. 666687-17), Washington University in St. Louis (no. 201102222), and the Nationwide Children’s Hospital (no. 2020N0022) approved the study in accordance with the Declaration of Helsinki. Written informed consent was obtained from participants aged 18 years and older and from the guardians of participants aged under 18 years. Written assent was obtained from participants aged 13 – 17 years and oral assent was obtained from younger children.

### Experimental design

Task-based iEEG data were derived from the encoding phase of two visual memory recognition tasks that have been used extensively to study memory in adults and children across neuroimaging modalities, including iEEG ^33^. In the blocked-trial paradigm, participants encode a set of 40 indoor and outdoor scenes and classify each as indoor/outdoor in preparation for a self-paced old/new recognition test of all 40 studied scenes intermixed with 20 new scenes as foils^2,3,9,13,45,50,76–79^. In the single trial paradigm, participants encode three shapes in a specific spatiotemporal sequence in preparation for a self-paced old/new recognition test of sequences that match exactly or mismatch on one dimension ^51,80–82^ (i.e., shape identity, spatial position, or temporal order; cf. ^83–88^). Both paradigms use visual stimuli to avoid potential confounds on memory with verbal material in children. The encoding phases of the two paradigms are similar because, in both paradigms, participants encode visual stimuli (3000ms, 500-1500ms inter-trial interval) in preparation for a self-paced, two-alternative forced choice recognition test. We ensured that on-task data reflected task engagement by only analyzing iEEG data during the viewing of stimuli that were attended during encoding, as indexed by a correct indoor/outdoor classification of each scene in the blocked-trial paradigm and correct old/new classification of each sequence in the single-trial paradigm ^9,13,50,51,79–81^. For a schematic of both visual memory tasks, see Figure S28. For task-free data, participants were instructed to sit quietly with their eyes open, fixating on the center of a computer monitor for five minutes. If no formal task-free task was administered, task-free data was taken from natural rest in continuous 24/7 iEEG recordings.

### Behavioral analysis

Both visual memory tasks test memory in a two-alternative forced choice design, permitting the use of similar measures of memory performance across tasks ^33^. For both tasks, for all participants, we calculated the hit rate (i.e., number of previously studied stimuli that were correctly recognized as old/match out of all studied stimuli) and false alarm rate (number of new stimuli presented that were incorrectly identified as old/match out of all new/mismatched stimuli). Performance accuracy was calculated as hit rate minus false alarm rate to equate measures across memory tasks and correct for differences in an individual’s tendency to respond old/match or new/mismatch, respectively. For a summary of behavioral performance, see Figure 1C.

### iEEG acquisition and pre-processing

iEEG data were recorded at a sampling rate of 200-5000 Hz using Nihon Kohden JE120 Neurofax or Natus Quantum LTM recording systems, which at three sites were interfaced with the BCI2000 software. Data acquired >1000 Hz were resampled to 1000 Hz after the fact. As described below, spectral analysis was performed up to 60 Hz. Thus, the lowest sampling rate of 200 is well over the minimum Nyquist frequency required for analysis (i.e., 2 cycles/frequency = 120 Hz). For consistency, all data from both visual memory tasks and from task-free recordings were pre-processed using the same procedures. Raw electrophysiological data were filtered with 0.1-Hz high-pass and 300-Hz low-pass finite impulse response filters, and 60-Hz line noise harmonics were removed using a discrete Fourier transform. Task-based continuous data were demeaned and epoched into 3s trials (i.e., 0-3s from scene or study sequence onset). Continuous task-free data were also demeaned and transformed into 3s epochs with 25% overlap. All epoched data were manually inspected blind to electrode locations and experimental task parameters. Electrodes overlying seizure onset zones and electrodes and epochs displaying epileptiform activity or artifactual signal (from poor contact, machine noise, etc.) were excluded (mean proportion of rejected epochs = 16.96%, *SD* = 12.51). We employed a bipolar re-referencing strategy, which has been shown to minimize the impact of impedance and electrode size differences between sEEG and ECoG and thus maximize standardization across these two types of recordings ^89^. Neighboring electrodes within the same anatomical structure were re-referenced using consistent conventions (ECoG, anterior – posterior; sEEG, deep – surface). For ECoG grids, electrodes were referenced to neighboring electrodes on a row-by-row basis. An electrode was discarded if it did not have an adjacent neighbor, its neighbor was in a different anatomical structure, or both it and its neighbor were in white matter. Bipolar referencing yielded virtual channels that were located midway between the original physical electrodes. Data were then manually re-inspected to reject any trials with residual noise. Pre-processing routines used functions from the FieldTrip toolbox for MATLAB ^90^. All results were based on analysis of non-pathologic, artifact-free channels, ensuring that data represented healthy cortical tissue ^20^.

### Peak frequency detection

The irregular-resampling auto-spectral analysis (IRASA) was used to isolate peak oscillatory components from the aperiodic 1/ƒ slope ^91^ as implemented in FieldTrip ^90^. Peak detection was performed from 1 – 60 Hz, with 60 Hz line noise removed using a discrete Fourier Transformation. The IRASA method compresses and expands the epoched data with non-integer resampling factors to redistribute oscillatory components while leaving the 1/ƒ distribution intact. For each original and resampled data trace, the auto-spectrum was calculated using the fast Fourier transform after applying a Hanning window. The median was taken from the resampled auto-spectra to obtain the 1/ƒ component for each channel and subtracted from the original power spectrum to isolate oscillatory residuals. Peak detection was performed on the oscillatory residuals within the theta range (1.5 – 9 Hz) using a minimum prominence threshold of 0.5 Hz. For cases with a single identified peak, a 4.5 Hz threshold was used to classify the oscillation as slow or fast theta ^9,51^. When two peaks were identified, the lower-frequency peak was classified as slow theta and the higher-frequency peak as fast theta. When more than two peaks were detected within a sub-band, the peak with the greatest prominence was retained.

### iEEG localization

Macro-electrodes were surgically implanted for extra-operative recording based solely on clinical need. The electrodes were subdural electrode grids or strips with 10 mm spacing or stereoelectroencephalography electrodes with 3-10 mm spacing. Anatomical locations were determined by co-registering post-implantation computed tomography coordinates to pre-operative magnetic resonance (MR) images, as implemented in FieldTrip ^90^, FreeSurfer ^92^, iELVis ^93^ or VERA^94^. Electrode locations were then projected into standard MNI space and bipolar channel locations (see preprocessing) were projected at the midpoint between their contributing electrodes. Based on these MNI coordinates, the *R* package *label4MRI* v1.2 (https://github.com/yunshiuan/label4MRI) was used to categorize each channel into its corresponding Brodmann area, which were then grouped according to the DKT atlas ^95^.

### Structural imaging and regional gray matter volume

T1-weighted MRI scans were acquired as part of routine preoperative procedures. Parcellation of cortex into regions of interest (ROI) was performed based on standard procedures implemented within FreeSurfer ^92^. Regional GMVs were then categorized based on the DKT atlas ^96^. GMV from each ROI was calculated using FreeSurfer ^92^. Volumes were calculated for left and right ROIs and averaged across hemispheres for analysis.

### Statistical analysis

Data were imported into *R* version 4.2.3 (R Core Team, 2020) with the aid of the *tidyverse* package ^97^ and analyzed using linear and nonlinear mixed-effects models fit by restricted maximum likelihood (REML) using *lme4* ^98^ and *splines* (R Core Team, 2020). *P*-values for region-specific models were estimated using the summary function from the *lmerTest* package, which is based on Satterthwaite’s degrees of freedom ^99^, and Type II Wald Tests from the *car* package ^100^ for examination of whole-brain effects (i.e., models which included all ROIs). Effects were plotted using the package gg*effects* ^101^ and *ggplot2* ^102^. Spearman correlations were used to assess structure-function relationships without the effect of age, with coefficients used to plot region-specific relationships between peak theta activities and GMV across the whole brain. Statistical significance was adjusted using the False Discovery Rate with an alpha threshold of .05. Task was entered as an unordered factor using sum-to-zero contrast coding and age was specified as a continuous predictor. See Table 1 below for a summary of the main analyses, including the types of models employed and their fixed and random effects structures.

**Table 1.** Summary of main statistical tests including number of models computed, outcome variables, fixed effects, and random effects.

| Analysis | #Models | Type | Outcome | Fixed Effects | Random Effects | FDR |
| --- | --- | --- | --- | --- | --- | --- |
| Association vs sensorimotor | 1 | Nonlinear mixed effects regression | Theta Frequency | 1. Age (continuous; one spline, 2 knots)<br>2. Cortex type (categorical) | 1. Participant (intercept)<br>2. ROI (intercept)<br>3. Channel (nested under participant) | No |
| Attentional state | 21 | Linear mixed effects regression | Theta Frequency | 1. Age (continuous)<br>2. Condition (categorical) | 1. Participant (intercept)<br>2. Channel (nested under participant)<br>3. Task (intercept) | Yes |
| Memory (task-based) | 21 | General linear regression | Accuracy | 1. Age (continuous)<br>2. Frequency (continuous) | NA | Yes |
| Memory (task-free) | 21 | General linear regression | Accuracy | 1. Age (continuous)<br>2. Theta Frequency (continuous) | NA | Yes |
| Gray matter volume (task-based) | 20 | Linear mixed effects regression | Theta Frequency | 1. Age (continuous)<br>2. GMV (continuous)<br>3. Intracranial volume (continuous) | 1. Participant (intercept)<br>2. Channel (nested under participant)<br>3. Task (intercept) | Yes |
| Gray matter volume (task-free) | 20 | Linear mixed effects regression | Theta Frequency | 1. Age (continuous)<br>2. GMV (continuous)<br>3. Intracranial volume (continuous) | 1. Participant (intercept)<br>2. Channel (nested under participant)<br>3. Task (intercept) | Yes |

*Note:* #Models refers to the number of models performed for each analysis; outcome = dependent variable; Cortex type has two levels of association and sensorimotor; Condition has two levels of task-free and task-based; False Discovery Rate (FDR) corrections were applied to all Analyses apart from “Association vs sensorimotor”, given that one model was computed.

In our preregistration, we specified that we would apply a spline to age to model potential non-linear effects of age on slow and fast theta frequencies for each ROI, as well as a random effect of task-free recording type (eyes open vs eyes closed). However, in doing so, models indicated nonconvergence. To reduce model complexity, we modeled age as a linear predictor and removed recording type as a random effect in our analysis of each ROI ^33^. For analyses testing hypotheses (a) and (b), where we tested differences in MTL and association and sensorimotor cortices, we had sufficient power to model nonlinear differences. Specifically, we used linear mixed-effects models with a single spline with two internal knots on age specified using the *splines* package. The model uses a natural spline with two degrees of freedom, which corresponds to a single spline with two knots. This spline approach allows for a nonlinear relationship between age and theta frequencies by dividing the age range into three segments. The choice of two knots reflects a balance between flexibility and model complexity, ensuring that we can model age-related differences without introducing excessive parameters. This approach enables the model to capture potential nonlinear patterns in the data, which is necessary to test our hypotheses.

Also note that when contrast coding is explicitly described, the need for post-hoc testing is eliminated (for a detailed discussion of contrast coding in linear mixed-effects regressions, please see ^103^). Further, for modeled effects, an 83% confidence interval (CI) threshold was used given that this approach corresponds to the 5% significance level with non-overlapping estimates ^104,105^. In order to isolate outliers for variables that were specified as outcomes (i.e., theta frequencies, memory performance), we used Tukey’s method ^106^, which identifies outliers as exceeding ± 1.5 × inter-quartile range. The packages *ggseg* ^107^ and *ggsegDKT* were used to generate cortical plots based on DKT atlas nomenclature. Hypotheses (a) and (b) were tested using the following formula (note that in all models, asterisks (*) denote interaction terms, while + denotes additive terms. *B*_0_ denotes the intercept of the model, while *B*_1_, *B*_2_ etc. denote the chronological specification of fixed effect parameters):

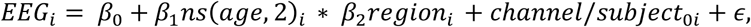

where *EEG* is slow or fast peak theta frequencies; *age* is age in years modeled with one spline term with two internal knots, and *region* encodes association and sensorimotor cortices; *channel* encodes region-specific channels nested under the random intercept of *participant*, and *participant* is the random intercept term of participant ID. To test hypothesis c, we employed the following model equation on a region-by-region basis:

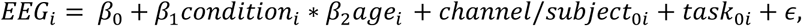

where *EEG* is peak slow or fast theta frequencies; *condition* encodes task-based and task-free recordings, age is age in years as a linear predictor; *channel* encodes region-specific channels nested under *participant*, and *participant* is participant ID, while *task* is a random intercept encoding whether the recording is derived from the working memory or scene recognition tasks.

Our exploratory analyses focused on relationships between GMV, behavioral performance, and slow or fast peak theta frequencies derived from task-based and task-free recordings. Here, our primary exploratory research questions were whether:

i. regional age-related variability in task-based and task-free slow and fast peak theta frequencies predict variability in memory performance, and;
ii. regional age-related variability in GMV predicts regional variability in task-based and task-free slow and fast peak theta frequencies.

These exploratory analyses were examined with general linear models with the following formulae:

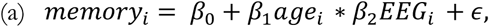

where *memory* is performance on the visual memory task(s), *age* is age in years, and *EEG* is slow or fast peak theta frequencies from each ROI and task-state (task-free, task-based).

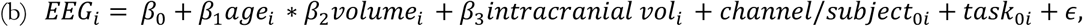

Here, *EEG* is task-based or task-free slow or fast peak theta frequencies; *age* is age in years; *volume* is regional GMV in mm^3^; *intracranial vol* refers to total intracranial volume, which was modelled as a covariate; *channel* encodes ROI-specific channels; *participant* is participant ID; *task* encodes whether the task recording was from the scene recognition or working memory task. As with the other models, each ROI was applied to the model equation described above. *Participant* was modeled as a random effect on the intercept, while *channel* was nested under participant. *Task* was also specified as a random effect on the intercept.

## Acknowledgements

This research was supported in part through the computational resources and staff contributions provided for the Quest high-performance computing facility at Northwestern University which is jointly supported by the Office of the Provost, the Office for Research, and Northwestern University Information Technology. We thank Dr. Kurtis I. Auguste, Dr. Joshua M. Rosenow, Professor Jack J. Lin and Professor Edward F. Chang for assistance with patient recruitment. Funding: R00NS115918 (E.L.J.), R01MH107512 (N.O.), R01NS21135 (R.T.K.), R00MH117226, P30AG013854, DGE-2234667 (Y.M.R.), T32MH067564 (Y.M.R. and C.C.), T32NS047987 (A.M.H.), P41EB018783. The funders had no role in study design, data collection and analysis, decision to publish or preparation of the manuscript.

## Author Contributions

Z.R.C., N.O., and E.L.J. designed the study. S.M.G., A.J.O.D., Y.M.R., C.R., A.M.H., E.A., O.K.M., S.S., I.S., F.G., D.K.S., P.B.W., K.D.L., S.U.S., J.Y.W., S.K.L., J.S.R., A.S., P.B., J.L.R., R.M.B., N.O., and E.L.J. recruited patients and/or collected data. Z.R.C., S.M.G., Q.Y., P.V., E.M.B.R., C.C., A.M.H., R.T.K., N.O., and E.L.J. preprocessed data. Z.R.C. and A.J.O.D. analyzed data. Z.R.C. visualized results. Z.R.C. and E.L.J. interpreted data. Z.R.C. drafted the manuscript. Z.R.C. and E.L.J. revised the manuscript. E.L.J. supervised the study. All authors provided feedback on the completed manuscript.

## References

1. Seger, S. E., Kriegel, J. L. S., Lega, B. C. & Ekstrom, A. D. Memory-related processing is the primary driver of human hippocampal theta oscillations. Neuron 111, 3119–3130.e4 (2023).

2. Yin, Q., Johnson, E. L. & Ofen, N. Neurophysiological mechanisms of cognition in the developing brain: Insights from intracranial EEG studies. Dev. Cogn. Neurosci. 64, 101312 (2023).

3. Ofen, N., Tang, L., Yu, Q. & Johnson, E. L. Memory and the developing brain: From description to explanation with innovation in methods. Dev. Cogn. Neurosci. 36, 100613 (2019).

4. Buzsáki, G. Theta Oscillations in the Hippocampus. Neuron 33, 325–340 (2002).

5. Backus, A. R., Schoffelen, J.-M., Szebényi, S., Hanslmayr, S. & Doeller, C. F. Hippocampal-Prefrontal Theta Oscillations Support Memory Integration. Curr. Biol. 26, 450–457 (2016).

6. Başar, E. & Güntekin, B. A review of brain oscillations in cognitive disorders and the role of neurotransmitters. Brain Res. 1235, 172–193 (2008).

7. Buzsáki, G. & Schomburg, E. W. What does gamma coherence tell us about inter-regional neural communication? Nat. Neurosci. 18, 484–489 (2015).

8. Helfrich, R. F. & Knight, R. T. Oscillatory Dynamics of Prefrontal Cognitive Control. Trends Cogn. Sci. 20, 916–930 (2016).

9. Johnson, E. L. et al. Dissociable oscillatory theta signatures of memory formation in the developing brain. Curr. Biol. 32, 1457–1469.e4 (2022).

10. Freschl, J., Al Azizi, L., Balboa, L., Kaldy, Z. & Blaser, E. The development of peak alpha frequency from infancy to adolescence and its role in visual temporal processing: A meta-analysis. Dev. Cogn. Neurosci. 101146 (2022).

11. Thuwal, K., Banerjee, A. & Roy, D. Aperiodic and periodic components of ongoing oscillatory brain dynamics link distinct functional aspects of cognition across adult lifespan. Eneuro 8, (2021).

12. Miles, J. T., Weaver, K. E., Webb, S. J. & Ojemann, J. G. Developmental relationships between the human alpha rhythm and intrinsic neural timescales are dependent on neural hierarchy. J. Neurophysiol. 135, 143–152 (2026).

13. Yin, Q. et al. Direct brain recordings reveal occipital cortex involvement in memory development. Neuropsychologia 148, 107625 (2020).

14. Musall, S., von Pföstl, V., Rauch, A., Logothetis, N. K. & Whittingstall, K. Effects of Neural Synchrony on Surface EEG. Cereb. Cortex 24, 1045–1053 (2014).

15. Palva, J. M. et al. Ghost interactions in MEG/EEG source space: A note of caution on inter-areal coupling measures. NeuroImage 173, 632–643 (2018).

16. Johnson, E. L., Kam, J. W., Tzovara, A. & Knight, R. T. Insights into human cognition from intracranial EEG: a review of audition, memory, internal cognition, and causality. J. Neural Eng. 17, 051001 (2020).

17. Johnson, E. L. & Knight, R. T. Intracranial recordings and human memory. Curr. Opin. Neurobiol. 31, 18–25 (2015).

18. Parvizi, J. & Kastner, S. Promises and limitations of human intracranial electroencephalography. Nat. Neurosci. 21, 474–483 (2018).

19. Liu, S. & Parvizi, J. Cognitive refractory state caused by spontaneous epileptic high-frequency oscillations in the human brain. Sci. Transl. Med. 11, eaax7830 (2019).

20. Rossini, L. et al. Seizure activity per se does not induce tissue damage markers in human neocortical focal epilepsy. Ann. Neurol. 82, 331–341 (2017).

21. Herweg, N. A., Solomon, E. A. & Kahana, M. J. Theta Oscillations in Human Memory. Trends Cogn. Sci. 24, 208–227 (2020).

22. Dede, A. J. O. et al. Episodic memory involves transient and sparse connectivity aligned to both internal and external events. PLOS Biol. 23, e3003481 (2025).

23. Llorens, A., et al. Decision and response monitoring during working memory are sequentially represented in the human insula. iScience 26, (2023).

24. Overton, J. A. et al. Distributed Intracranial Activity Underlying Human Decision-making Behavior. J. Neurosci. 45, e0572242024 (2025).

25. Johnson, E. L. et al. A rapid theta network mechanism for flexible information encoding. Nat. Commun. 14, 2872 (2023).

26. ter Wal, M., et al. Theta rhythmicity governs human behavior and hippocampal signals during memory-dependent tasks. Nat. Commun. 12, 7048 (2021).

27. Chen, D. et al. Hexadirectional Modulation of Theta Power in Human Entorhinal Cortex during Spatial Navigation. Curr. Biol. 28, 3310–3315.e4 (2018).

28. Kunz, L. et al. Hippocampal theta phases organize the reactivation of large-scale electrophysiological representations during goal-directed navigation. Sci. Adv. 5, eaav8192.

29. Begus, K. & Bonawitz, E. The rhythm of learning: Theta oscillations as an index of active learning in infancy. Dev. Cogn. Neurosci. 45, 100810 (2020).

30. Cellier, D., Riddle, J., Petersen, I. & Hwang, K. The development of theta and alpha neural oscillations from ages 3 to 24 years. Dev. Cogn. Neurosci. 50, 100969 (2021).

31. van Noordt, S., Heffer, T. & Willoughby, T. A developmental examination of medial frontal theta dynamics and inhibitory control. NeuroImage 246, 118765 (2022).

32. Schneider, J. M., Abel, A. D., Ogiela, D. A., Middleton, A. E. & Maguire, M. J. Developmental differences in beta and theta power during sentence processing. Dev. Cogn. Neurosci. 19, 19–30 (2016).

33. Cross, Z. R. et al. The development of aperiodic neural activity in the human brain. Nat. Hum. Behav. 10.1038/s41562-025-02270-x (2025) doi:10.1038/s41562-025-02270-x.

34. Doval, S. et al. When Maturation is Not Linear: Brain Oscillatory Activity in the Process of Aging as Measured by Electrophysiology. Brain Topogr. 10.1007/s10548-024-01064-0 (2024) doi:10.1007/s10548-024-01064-0.

35. Overbye, K., Huster, R. J., Walhovd, K. B., Fjell, A. M. & Tamnes, C. K. Development of the P300 from childhood to adulthood: a multimodal EEG and MRI study. Brain Struct. Funct. 223, 4337–4349 (2018).

36. Sui, J., Huster, R., Yu, Q., Segall, J. & Calhoun, V. Function-structure associations of the brain: evidence from multimodal connectivity and covariance studies. Neuroimage 102P1, 11–23. (2014).

37. Whitford, T. J. et al. Brain maturation in adolescence: Concurrent changes in neuroanatomy and neurophysiology. Hum. Brain Mapp. 28, 228–237 (2007).

38. Bethlehem, R. A. I. et al. Brain charts for the human lifespan. Nature 604, 525–533 (2022).

39. Gogtay, N. et al. Dynamic mapping of human cortical development during childhood through early adulthood. Proc. Natl. Acad. Sci. 101, 8174–8179 (2004).

40. Groeschel, S., Vollmer, B., King, M. & Connelly, A. Developmental changes in cerebral grey and white matter volume from infancy to adulthood. Int. J. Dev. Neurosci. 28, 481–489 (2010).

41. Grydeland, H. et al. Waves of maturation and senescence in micro-structural MRI markers of human cortical myelination over the lifespan. Cereb. Cortex 29, 1369–1381 (2019).

42. Hill, J. et al. Similar patterns of cortical expansion during human development and evolution. Proc. Natl. Acad. Sci. 107, 13135–13140 (2010).

43. Sydnor, V. J. et al. Neurodevelopment of the association cortices: Patterns, mechanisms, and implications for psychopathology. Neuron 109, 2820–2846 (2021).

44. Gogtay, N. et al. Dynamic mapping of normal human hippocampal development. Hippocampus 16, 664–672 (2006).

45. Ofen, N. et al. Development of the declarative memory system in the human brain. Nat. Neurosci. 10, 1198–1205 (2007).

46. Wilke, M., Krägeloh-Mann, I. & Holland, S. K. Global and local development of gray and white matter volume in normal children and adolescents. Exp. Brain Res. 178, 296–307 (2007).

47. Hill, P. F., King, D. R., Lega, B. C. & Rugg, M. D. Comparison of fMRI correlates of successful episodic memory encoding in temporal lobe epilepsy patients and healthy controls. NeuroImage 207, 116397 (2020).

48. Johnson, E. L. & Knight, R. T. How Can iEEG Be Used to Study Inter-Individual and Developmental Differences? in Intracranial EEG: A Guide for Cognitive Neuroscientists 143–154 (Springer, 2023).

49. Desikan, R. S. et al. An automated labeling system for subdividing the human cerebral cortex on MRI scans into gyral based regions of interest. Neuroimage 31, 968–980 (2006).

50. Johnson, E. L., Tang, L., Yin, Q., Asano, E. & Ofen, N. Direct brain recordings reveal prefrontal cortex dynamics of memory development. Sci. Adv. 4, eaat3702 (2018).

51. Gray, S. M. et al. Opposing effects of slow and fast theta synchrony on working memory in the human hippocampal-orbitofrontal network. bioRxiv 2026–05 (2026).

52. Gao, W. A hierarchical model of early brain functional network development. Trends Cogn. Sci. 29, 855–868 (2025).

53. Casey, B., Giedd, J. N. & Thomas, K. M. Structural and functional brain development and its relation to cognitive development. Biol. Psychol. 54, 241–257 (2000).

54. Wang, D., Parish, G., Shapiro, K. L. & Hanslmayr, S. Interaction between Theta Phase and Spike Timing-Dependent Plasticity Simulates Theta-Induced Memory Effects. eneuro 10, ENEURO.0333-22.2023 (2023).

55. Siegle, J. H. & Wilson, M. A. Enhancement of encoding and retrieval functions through theta phase-specific manipulation of hippocampus. eLife 3, e03061 (2014).

56. Skaggs, W. E., McNaughton, B. L., Wilson, M. A. & Barnes, C. A. Theta phase precession in hippocampal neuronal populations and the compression of temporal sequences. Hippocampus 6, 149– 172 (1996).

57. Bakker, I., Takashima, A., Van Hell, J. G., Janzen, G. & McQueen, J. M. Changes in theta and beta oscillations as signatures of novel word consolidation. J. Cogn. Neurosci. 27, 1286–1297 (2015).

58. Klimesch, W. EEG alpha and theta oscillations reflect cognitive and memory performance: a review and analysis. Brain Res. Rev. 29, 169–195 (1999).

59. Tan, E. et al. Theta activity and cognitive functioning: Integrating evidence from resting-state and task-related developmental electroencephalography (EEG) research. Dev. Cogn. Neurosci. 67, 101404 (2024).

60. Jacobs, J., Hwang, G., Curran, T. & Kahana, M. J. EEG oscillations and recognition memory: Theta correlates of memory retrieval and decision making. NeuroImage 32, 978–987 (2006).

61. Soutschek, A., Moisa, M., Ruff, C. C. & Tobler, P. N. Frontopolar theta oscillations link metacognition with prospective decision making. Nat. Commun. 12, 3943 (2021).

62. Crivelli-Decker, J., Hsieh, L.-T., Clarke, A. & Ranganath, C. Theta oscillations promote temporal sequence learning. Neurobiol. Learn. Mem. 153, 92–103 (2018).

63. Sun, L. & Bao, L. Neuronal theta oscillation of hippocampal ensemble and memory function. Behav. Brain Res. 481, 115429 (2025).

64. Bitzenhofer, S. H., Sieben, K., Siebert, K. D., Spehr, M. & Hanganu-Opatz, I. L. Oscillatory Activity in Developing Prefrontal Networks Results from Theta-Gamma-Modulated Synaptic Inputs. Cell Rep. 11, 486–497 (2015).

65. Leung, L. S. & Law, C. S. Phasic modulation of hippocampal synaptic plasticity by theta rhythm. Behav. Neurosci. 134, 595 (2020).

66. Mizuseki, K., Sirota, A., Pastalkova, E. & Buzsáki, G. Theta oscillations provide temporal windows for local circuit computation in the entorhinal-hippocampal loop. Neuron 64, 267–280 (2009).

67. Constable, R. T. Structure–function relationships across scales: implications for atlases. Front. Neurosci. 19, 1750272 (2025).

68. Suárez, L. E., Markello, R. D., Betzel, R. F. & Misic, B. Linking Structure and Function in Macroscale Brain Networks. Trends Cogn. Sci. 24, 302–315 (2020).

69. Ghetti, S. & Bunge, S. A. Neural changes underlying the development of episodic memory during middle childhood. Dev. Cogn. Neurosci. 2, 381–395 (2012).

70. Herbet, G. & Duffau, H. Revisiting the Functional Anatomy of the Human Brain: Toward a Meta-Networking Theory of Cerebral Functions. Physiol. Rev. 100, 1181–1228 (2020).

71. Arns, M., Conners, C. K. & Kraemer, H. C. A decade of EEG theta/beta ratio research in ADHD: a meta-analysis. J. Atten. Disord. 17, 374–383 (2013).

72. Sonkusare, S., Breakspear, M. & Guo, C. Naturalistic Stimuli in Neuroscience: Critically Acclaimed. Trends Cogn. Sci. 23, 699–714 (2019).

73. Campos, A., Loyola-Navarro, R., González, C. & Iverson, P. Resting-state electroencephalogram and speech perception in young children with developmental language disorder. Brain Sci. 15, 219 (2025).

74. Musaeus, C. S. et al. EEG theta power is an early marker of cognitive decline in dementia due to Alzheimer’s disease. J. Alzheimer’s Dis. 64, 1359–1371 (2018).

75. Perez, V., Duque, A., Hidalgo, V. & Salvador, A. EEG frequency bands in subjective cognitive decline: A systematic review of resting state studies. Biol. Psychol. 191, 108823 (2024).

76. Chai, X. J., Ofen, N., Jacobs, L. F. & Gabrieli, J. D. Scene complexity: influence on perception, memory, and development in the medial temporal lobe. Front. Hum. Neurosci. 4, 1021 (2010).

77. Ofen, N., Chai, X. J., Schuil, K. D., Whitfield-Gabrieli, S. & Gabrieli, J. D. The development of brain systems associated with successful memory retrieval of scenes. J. Neurosci. 32, 10012–10020 (2012).

78. Tang, L., Shafer, A. T. & Ofen, N. Prefrontal Cortex Contributions to the Development of Memory Formation. Cereb. Cortex 28, 3295–3308 (2018).

79. Yin, Q. et al. Distinct neurophysiological features and memory representations along the long axis of the developing medial temporal lobe. bioRxiv 2025–10 (2025).

80. Yarbrough, J. B., Shi, L., Chattopadhyay, K., Knight, R. T. & Johnson, E. L. One of these things is not like the others: Theta, beta, & ERP dynamics of mismatch detection. bioRxiv 2025–07 (2025).

81. Shi, L. et al. Distributed theta networks support the control of working memory: Evidence from scalp and intracranial EEG. bioRxiv 2025–08 (2025).

82. Ack, S. E. et al. Differential roles of frontoparietal regions in working memory and decision-making. bioRxiv 2026.09.23.753872 (2026) doi:10.64898/2026.09.23.753872.

83. Davoudi, S., Parto Dezfouli, M., Knight, R. T., Daliri, M. R. & Johnson, E. L. Prefrontal lesions disrupt posterior alpha–gamma coordination of visual working memory representations. J. Cogn. Neurosci. 33, 1798–1810 (2021).

84. Dezfouli, M. P., Davoudi, S., Knight, R. T., Daliri, M. R. & Johnson, E. L. Prefrontal lesions disrupt oscillatory signatures of spatiotemporal integration in working memory. Cortex 138, 113–126 (2021).

85. Johnson, E. L. et al. Dynamic frontotemporal systems process space and time in working memory. PLoS Biol. 16, e2004274 (2018).

86. Johnson, E. L. et al. Orbitofrontal cortex governs working memory for temporal order. Curr. Biol. 32, R410–R411 (2022).

87. Johnson, E. L. et al. Bidirectional frontoparietal oscillatory systems support working memory. Curr. Biol. 27, 1829–1835 (2017).

88. Johnson, E. L. et al. Spectral imprints of working memory for everyday associations in the frontoparietal network. Front. Syst. Neurosci. 12, 65 (2019).

89. Mercier, M. R. et al. Advances in human intracranial electroencephalography research, guidelines and good practices. Neuroimage 260, 119438 (2022).

90. Oostenveld, R., Fries, P., Maris, E. & Schoffelen, J.-M. FieldTrip: open source software for advanced analysis of MEG, EEG, and invasive electrophysiological data. Comput. Intell. Neurosci. 2011, 1–9 (2011).

91. Wen, H. & Liu, Z. Separating fractal and oscillatory components in the power spectrum of neurophysiological signal. Brain Topogr. 29, 13–26 (2016).

92. Fischl, B. FreeSurfer. Neuroimage 62, 774–781 (2012).

93. Groppe, D. M. et al. iELVis: An open source MATLAB toolbox for localizing and visualizing human intracranial electrode data. J. Neurosci. Methods 281, 40–48 (2017).

94. Adamek, M., Swift, J. & Brunner, P. VERA-Versatile electrode localization Framework. Prepr. Version Doi-Release 10, (2022).

95. Schmidt, F. et al. Age-related changes in “cortical” 1/f dynamics are linked to cardiac activity. 10.7554/elife.100605.1 (2024) doi:10.7554/elife.100605.1.

96. Klein, A. & Tourville, J. 101 Labeled Brain Images and a Consistent Human Cortical Labeling Protocol. Front. Neurosci. 6, (2012).

97. Wickham, H. et al. Welcome to the Tidyverse. J. Open Source Softw. 4, 1686 (2019).

98. Bates, D. M. lme4: Mixed-effects modeling with R. (2010).

99. Kuznetsova, A., Brockhoff, P. B. & Christensen, R. H. lmerTest package: tests in linear mixed effects models. J. Stat. Softw. 82, 1–26 (2017).

100. Fox, J. et al. The car package. R Found. Stat. Comput. 1109, 1431 (2007).

101. Lüdecke, D. ggeffects: Tidy data frames of marginal effects from regression models. J. Open Source Softw. 3, 772 (2018).

102. Wickham, H. & Wickham, H. Data analysis. Ggplot2 Elegant Graph. Data Anal. 189–201 (2016).

103. Brehm, L. & Alday, P. M. Contrast coding choices in a decade of mixed models. J. Mem. Lang. 125, 104334 (2022).

104. Austin, P. C. & Hux, J. E. A brief note on overlapping confidence intervals. J. Vasc. Surg. 36, 194–195 (2002).

105. MacGregor-Fors, I. & Payton, M. E. Contrasting diversity values: statistical inferences based on overlapping confidence intervals. PLoS One 8, e56794 (2013).

106. 106. Tukey, J. W. Exploratory Data Analysis. vol. 2 (Reading, MA, 1977).

107. Mowinckel, A. M. & Vidal-Piñeiro, D. Visualization of brain statistics with R packages ggseg and ggseg3d. Adv. Methods Pract. Psychol. Sci. 3, 466–483 (2020).

