## supplementary material for "The development of slow and fast theta oscillations in the human brain"

**S1.** Summary of key dependent and independent variables across each region of interest for task-based slow theta.

| Region | Condition | Age Range | | Memory Range | | Frequency Range | | GMV Range | | n Subject | n Channel |
|---|---|---|---|---|---|---|---|---|---|---|---|
| amygdala | Task | 11.17 | 41.0 | 0.16 | 1.00 | 1.71 | 5.62 | 1735.5 | 2164.75 | 14 | 36 |
| caudal anterior cingulate cortex | Task | 6.17 | 54.0 | 0.02 | 1.00 | 1.71 | 5.86 | 959.25 | 4544.5 | 36 | 151 |
| caudal middle frontal gyrus | Task | 5.94 | 41.0 | 0.01 | 1.00 | 1.71 | 6.59 | 8626.25 | 24964.25 | 68 | 767 |
| fusiform gyrus | Task | 6.17 | 38.0 | 0.02 | 1.00 | 1.71 | 7.08 | 2752.5 | 11793.5 | 52 | 282 |
| hippocampus | Task | 9.46 | 51.0 | 0.16 | 1.00 | 1.71 | 6.10 | 1022.25 | 2544 | 26 | 96 |
| inferior frontal gyrus | Task | 5.94 | 54.0 | 0.01 | 1.00 | 1.71 | 5.62 | 2268.83 | 5171.5 | 55 | 384 |
| inferior parietal cortex | Task | 5.94 | 51.0 | 0.01 | 0.96 | 1.71 | 6.59 | 6130 | 23277 | 56 | 459 |
| inferior temporal cortex | Task | 5.94 | 51.0 | 0.02 | 1.00 | 1.71 | 5.13 | 3638.5 | 17129.5 | 55 | 234 |
| insula | Task | 9.46 | 38.0 | 0.16 | 0.97 | 1.71 | 4.88 | 5374.5 | 7468.5 | 17 | 100 |
| lateral occipital cortex | Task | 6.17 | 28.1 | 0.02 | 1.00 | 1.71 | 7.08 | 4049.75 | 8889.12 | 40 | 269 |
| lateral orbitofrontal cortex | Task | 5.94 | 43.0 | 0.01 | 1.00 | 1.71 | 4.88 | 6668 | 12474.5 | 36 | 97 |
| medial orbitofrontal cortex | Task | 5.94 | 41.0 | 0.01 | 0.97 | 1.71 | 6.10 | 3649 | 6241.5 | 23 | 126 |
| middle temporal cortex | Task | 5.94 | 54.0 | 0.01 | 1.00 | 1.71 | 6.84 | 5741.5 | 22048 | 67 | 443 |
| parahippocampal gyrus | Task | 5.94 | 38.0 | 0.02 | 1.00 | 1.71 | 5.62 | 392.25 | 2579.75 | 44 | 142 |
| postcentral gyrus | Task | 5.94 | 32.0 | 0.01 | 0.96 | 1.71 | 5.62 | 6759.5 | 15504.5 | 48 | 195 |
| posterior cingulate cortex | Task | 6.17 | 51.0 | 0.02 | 1.00 | 1.71 | 6.10 | 1252 | 4043 | 32 | 116 |
| precentral gyrus | Task | 5.94 | 26.9 | 0.01 | 0.96 | 1.71 | 5.13 | 5302.25 | 11415 | 47 | 116 |
| rostral middle frontal gyrus | Task | 5.94 | 54.0 | 0.01 | 1.00 | 1.71 | 5.86 | 7832 | 17498 | 53 | 224 |
| subcortical | Task | 9.46 | 51.0 | 0.24 | 0.97 | 1.71 | 4.88 | NA | NA | 15 | 51 |
| superior parietal cortex | Task | 6.17 | 32.0 | 0.02 | 0.96 | 1.71 | 4.64 | 5766.66 | 15529.16 | 22 | 88 |
| superior temporal cortex | Task | 5.94 | 54.0 | 0.01 | 0.98 | 1.71 | 4.88 | 4813.25 | 12732.75 | 57 | 276 |

**S2.** Summary of key dependent and independent variables across each region of interest for task-free slow theta.

| Region | Condition | Age Range | | Frequency Range | | GMV Range | | n Subject | n Channel |
|---|---|---|---|---|---|---|---|---|---|
| amygdala | Rest | 16.84 | 34.84 | 1.709 | 3.1738 | 1296.25 | 1962.50 | 7 | 13 |
| caudal anterior cingulate cortex | Rest | 6.17 | 38.00 | 1.709 | 5.3711 | 959.25 | 4544.50 | 21 | 71 |
| caudal middle frontal gyrus | Rest | 5.94 | 45.00 | 1.709 | 7.0801 | 8753.50 | 24964.25 | 47 | 418 |
| fusiform gyrus | Rest | 6.17 | 54.00 | 1.709 | 5.127 | 2752.50 | 11793.50 | 44 | 227 |
| hippocampus | Rest | 9.46 | 32.00 | 1.709 | 6.3477 | 1482.75 | 2327.25 | 9 | 38 |
| inferior frontal gyrus | Rest | 5.94 | 54.00 | 1.709 | 5.8594 | 2268.83 | 5100.00 | 33 | 227 |
| inferior parietal cortex | Rest | 5.94 | 54.00 | 1.709 | 5.8594 | 10405.00 | 23277.00 | 41 | 240 |
| inferior temporal cortex | Rest | 5.94 | 54.00 | 1.709 | 6.1035 | 3638.50 | 17129.50 | 41 | 180 |
| insula | Rest | 9.46 | 54.00 | 1.709 | 5.6152 | 4852.00 | 6928.00 | 7 | 39 |
| lateral occipital cortex | Rest | 6.17 | 34.84 | 1.709 | 4.8828 | 4801.50 | 8889.13 | 33 | 155 |
| lateral orbitofrontal cortex | Rest | 5.94 | 45.00 | 1.709 | 4.3945 | 6849.00 | 12474.50 | 24 | 58 |
| medial orbitofrontal cortex | Rest | 5.94 | 32.00 | 1.709 | 4.3945 | 3843.00 | 6241.50 | 13 | 40 |
| middle temporal cortex | Rest | 5.94 | 54.00 | 1.709 | 6.3477 | 5741.50 | 22048.00 | 41 | 249 |
| parahippocampal gyrus | Rest | 5.94 | 54.00 | 1.709 | 4.6387 | 392.25 | 2352.50 | 32 | 108 |
| postcentral gyrus | Rest | 5.94 | 38.00 | 1.709 | 4.6387 | 6759.50 | 14982.50 | 34 | 119 |
| posterior cingulate cortex | Rest | 6.17 | 43.00 | 1.709 | 4.3945 | 1252.00 | 4043.00 | 22 | 62 |
| precentral gyrus | Rest | 5.94 | 38.00 | 1.709 | 7.3242 | 5302.25 | 11415.00 | 33 | 74 |
| rostral middle frontal gyrus | Rest | 5.94 | 38.00 | 1.709 | 5.3711 | 9210.50 | 17498.00 | 30 | 109 |
| subcortical | Rest | 9.46 | 43.00 | 1.709 | 6.1035 | NA | NA | 12 | 41 |
| superior parietal cortex | Rest | 6.17 | 43.00 | 1.709 | 5.8594 | 5766.67 | 15529.17 | 14 | 34 |
| superior temporal cortex | Rest | 5.94 | 43.00 | 1.709 | 6.3477 | 4951.50 | 12732.75 | 37 | 172 |

**S3.** Summary of key dependent and independent variables across each region of interest for task-based fast theta.

| Region | Condition | Age Range | | Memory Range | | Frequency Range | | GMV Range | | n Subject | n Channel |
|---|---|---|---|---|---|---|---|---|---|---|---|
| amygdala | Task | 11.17 | 41.00 | 0.16 | 1.00 | 3.91 | 8.54 | 1735.5 | 2164.75 | 14 | 36 |
| caudal anterior cingulate cortex | Task | 6.17 | 54.00 | 0.02 | 1.00 | 3.66 | 8.79 | 959.25 | 4544.5 | 36 | 151 |
| caudal middle frontal gyrus | Task | 5.94 | 41.00 | 0.01 | 1.00 | 3.42 | 8.79 | 8626.25 | 24964.25 | 68 | 767 |
| fusiform gyrus | Task | 6.17 | 38.00 | 0.02 | 1.00 | 3.17 | 8.79 | 2752.5 | 11793.5 | 52 | 282 |
| hippocampus | Task | 9.46 | 51.00 | 0.16 | 1.00 | 3.17 | 8.79 | 1022.25 | 2544 | 26 | 96 |
| inferior frontal gyrus | Task | 5.94 | 54.00 | 0.01 | 1.00 | 4.15 | 8.79 | 2268.83 | 5171.5 | 55 | 384 |
| inferior parietal cortex | Task | 5.94 | 51.00 | 0.01 | 0.96 | 2.93 | 8.79 | 6130 | 23277 | 56 | 459 |
| inferior temporal cortex | Task | 5.94 | 51.00 | 0.02 | 1.00 | 3.66 | 8.54 | 3638.5 | 17129.5 | 55 | 234 |
| insula | Task | 9.46 | 38.00 | 0.16 | 0.97 | 3.66 | 8.79 | 5374.5 | 7468.5 | 17 | 100 |
| lateral occipital cortex | Task | 6.17 | 28.11 | 0.02 | 1.00 | 3.42 | 8.79 | 4049.75 | 8889.12 | 40 | 269 |
| lateral orbitofrontal cortex | Task | 5.94 | 43.00 | 0.01 | 1.00 | 3.17 | 8.79 | 6668 | 12474.5 | 36 | 97 |
| medial orbitofrontal cortex | Task | 5.94 | 41.00 | 0.01 | 0.97 | 3.66 | 8.54 | 3649 | 6241.5 | 23 | 126 |
| middle temporal cortex | Task | 5.94 | 54.00 | 0.01 | 1.00 | 3.17 | 8.79 | 5741.5 | 22048 | 67 | 443 |
| parahippocampal gyrus | Task | 5.94 | 38.00 | 0.02 | 1.00 | 4.00 | 8.54 | 392.25 | 2579.75 | 44 | 142 |
| postcentral gyrus | Task | 5.94 | 32.00 | 0.01 | 0.96 | 3.91 | 8.79 | 6759.5 | 15504.5 | 48 | 195 |
| posterior cingulate cortex | Task | 6.17 | 51.00 | 0.02 | 1.00 | 3.91 | 8.79 | 1252 | 4043 | 32 | 116 |
| precentral gyrus | Task | 5.94 | 26.89 | 0.01 | 0.96 | 2.93 | 8.79 | 5302.25 | 11415 | 47 | 116 |
| rostral middle frontal gyrus | Task | 5.94 | 54.00 | 0.01 | 1.00 | 4.39 | 8.79 | 7832 | 17498 | 53 | 224 |
| subcortical | Task | 9.46 | 51.00 | 0.24 | 0.97 | 3.91 | 8.79 | NA | NA | 15 | 51 |
| superior parietal cortex | Task | 6.17 | 32.00 | 0.02 | 0.96 | 3.66 | 8.79 | 5766.66 | 15529.16 | 22 | 88 |
| superior temporal cortex | Task | 5.94 | 54.00 | 0.01 | 0.98 | 3.42 | 8.79 | 4813.25 | 12732.75 | 57 | 276 |

**S4.** Summary of key dependent and independent variables across each region of interest for task-free fast theta.

| Region | Condition | Age Range | | Frequency Range | | GMV Range | | n Subject | n Channel |
|---|---|---|---|---|---|---|---|---|---|
| amygdala | Rest | 16.84 | 34.84 | 4.15 | 8.06 | 1296.25 | 1962.50 | 7 | 13 |
| caudal anterior cingulate cortex | Rest | 6.17 | 38.00 | 4.64 | 8.54 | 959.25 | 4544.50 | 21 | 71 |
| caudal middle frontal gyrus | Rest | 5.94 | 45.00 | 3.91 | 8.79 | 8753.50 | 24964.25 | 47 | 418 |
| fusiform gyrus | Rest | 6.17 | 54.00 | 3.17 | 8.79 | 2752.50 | 11793.50 | 44 | 227 |
| hippocampus | Rest | 9.46 | 32.00 | 3.66 | 8.79 | 1482.75 | 2327.25 | 9 | 38 |
| inferior frontal gyrus | Rest | 5.94 | 54.00 | 4.15 | 8.79 | 2268.83 | 5100.00 | 33 | 227 |
| inferior parietal cortex | Rest | 5.94 | 54.00 | 3.42 | 8.79 | 10405.00 | 23277.00 | 41 | 240 |
| inferior temporal cortex | Rest | 5.94 | 54.00 | 3.17 | 8.79 | 3638.50 | 17129.50 | 41 | 180 |
| insula | Rest | 9.46 | 54.00 | 3.66 | 8.79 | 4852.00 | 6928.00 | 7 | 39 |
| lateral occipital cortex | Rest | 6.17 | 34.84 | 3.42 | 8.79 | 4801.50 | 8889.13 | 33 | 155 |
| lateral orbitofrontal cortex | Rest | 5.94 | 45.00 | 4.39 | 8.54 | 6849.00 | 12474.50 | 24 | 58 |
| medial orbitofrontal cortex | Rest | 5.94 | 32.00 | 5.13 | 8.79 | 3843.00 | 6241.50 | 13 | 40 |
| middle temporal cortex | Rest | 5.94 | 54.00 | 3.42 | 8.79 | 5741.50 | 22048.00 | 41 | 249 |
| parahippocampal gyrus | Rest | 5.94 | 54.00 | 4.15 | 8.79 | 392.25 | 2352.50 | 32 | 108 |
| postcentral gyrus | Rest | 5.94 | 38.00 | 4.64 | 8.79 | 6759.50 | 14982.50 | 34 | 119 |
| posterior cingulate cortex | Rest | 6.17 | 43.00 | 4.64 | 8.79 | 1252.00 | 4043.00 | 22 | 62 |
| precentral gyrus | Rest | 5.94 | 38.00 | 4.64 | 8.79 | 5302.25 | 11415.00 | 33 | 74 |
| rostral middle frontal gyrus | Rest | 5.94 | 38.00 | 4.64 | 8.79 | 9210.50 | 17498.00 | 30 | 109 |
| subcortical | Rest | 9.46 | 43.00 | 4.64 | 8.54 | NA | NA | 12 | 41 |
| superior parietal cortex | Rest | 6.17 | 43.00 | 4.15 | 8.06 | 5766.67 | 15529.17 | 14 | 34 |
| superior temporal cortex | Rest | 5.94 | 43.00 | 4.39 | 8.79 | 4951.50 | 12732.75 | 37 | 172 |

**S5.** Summary of association and sensorimotor regions.

| **Association** | **Sensorimotor** |
|---|---|
| Caudal Anterior Cingulate | Lateral Occipital Cortex |
| Caudal Middle Frontal Gyrus | Precentral Gyrus |
| Fusiform Gyrus | Postcentral Gyrus |
| Inferior Frontal Gyrus | |
| Inferior Parietal Cortex | |
| Inferior Temporal Cortex | |
| Insula | |
| Lateral Orbitofrontal Cortex | |
| Medial Orbitofrontal Cortex | |
| Middle Temporal Cortex | |
| Parahippocampal Gyrus | |
| Posterior Cingulate Cortex | |
| Rostral Middle Frontal Gyrus | |
| Superior Parietal Cortex | |
| Superior Temporal Cortex | |

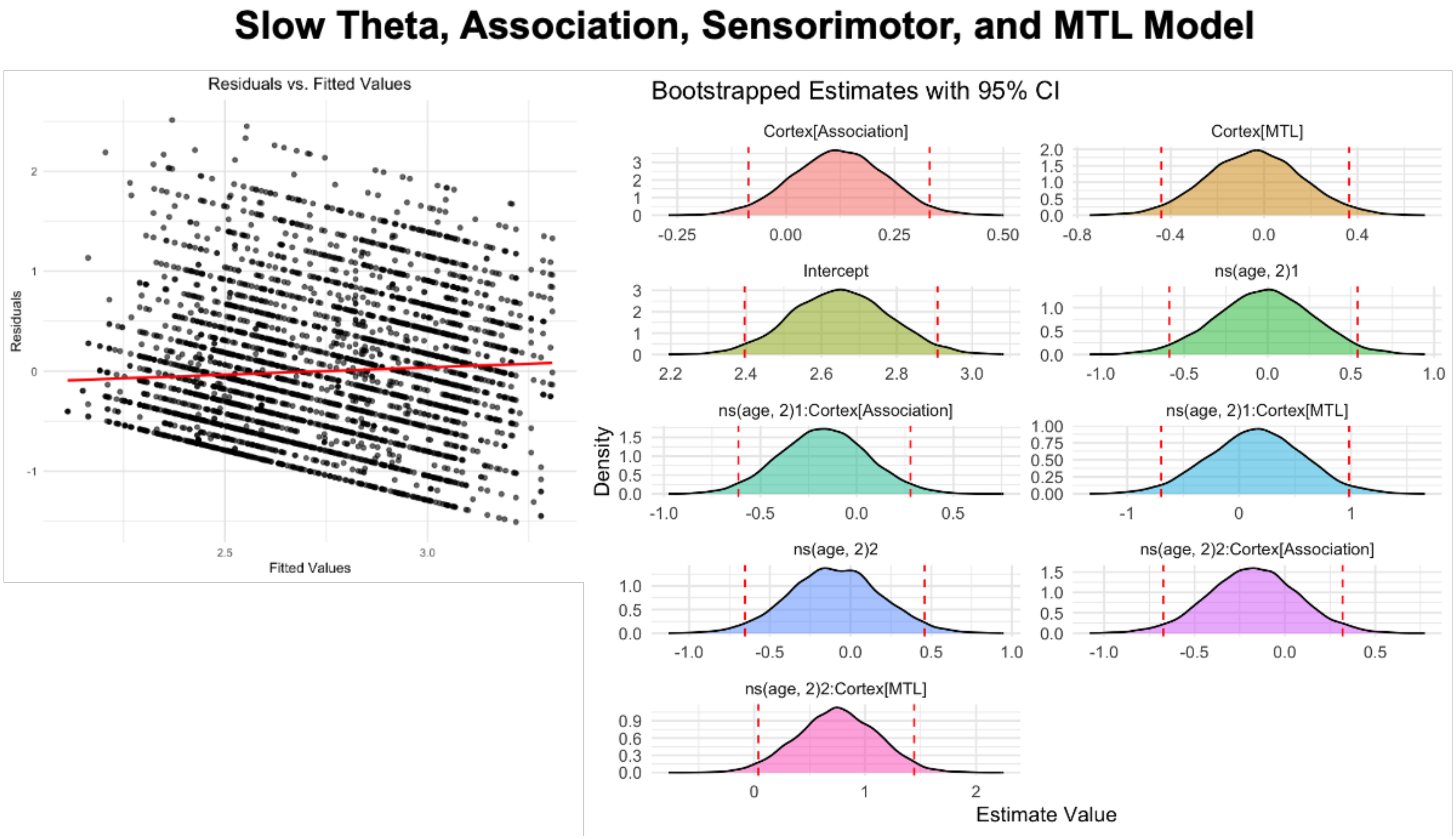


**S6. Residuals and bootstrapped beta coefficients for slow theta and association, sensorimotor and medial temporal lobe model.** Left: Q-Q plot of residuals of the mixed-effects model. Right: Bootstrapped confidence intervals for the intercept, main effects and interactions. Y-axis represents the density of the bootstrapped coefficients, and the x-axis represents the estimated beta values. The dashed red lines indicate the 95% confidence interval from the original mixed-model estimates.

### Fast Theta, Association, Sensorimotor, and MTL Model

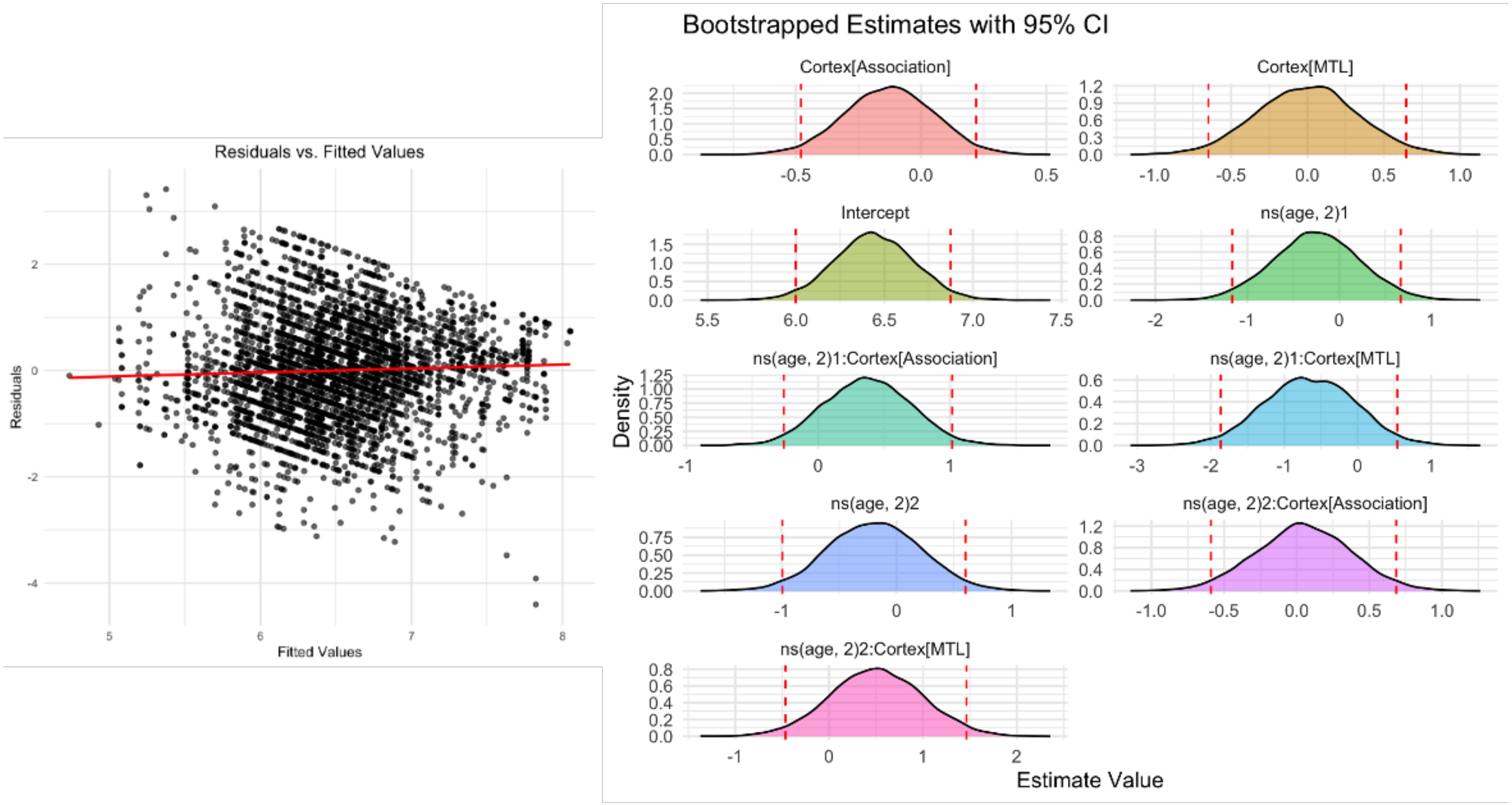


**S7. Residuals and bootstrapped beta coefficients for fast theta and association, sensorimotor and medial temporal lobe model.** Left: Q-Q plot of residuals of the mixed-effects model. Right: Bootstrapped confidence intervals for the intercept, main effects and interactions. Y-axis represents the density of the bootstrapped coefficients, and the x-axis represents the estimated beta values. The dashed red lines indicate the 95% confidence interval from the original mixed-model estimates.

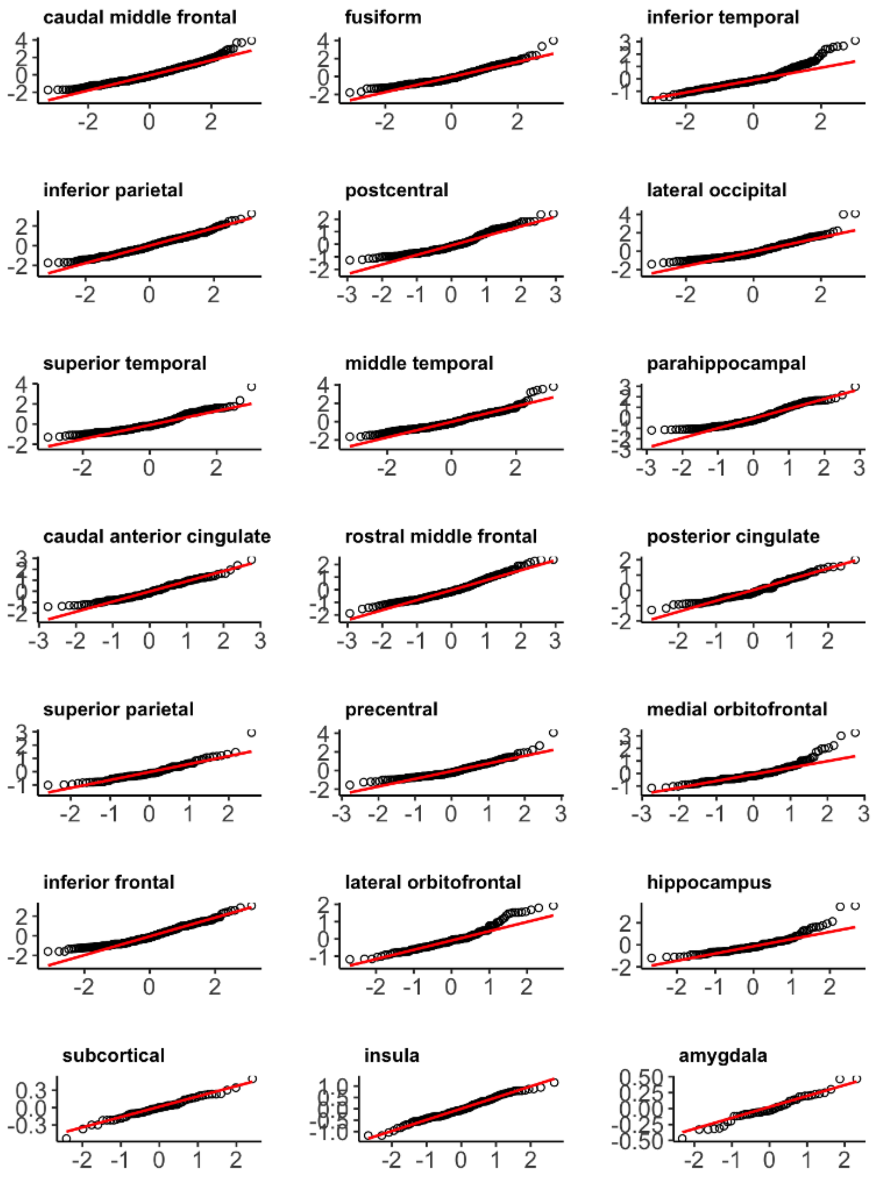


**S8.** Q-Q Plots of residuals for models examining age and attentional state on peak slow theta for each ROI.

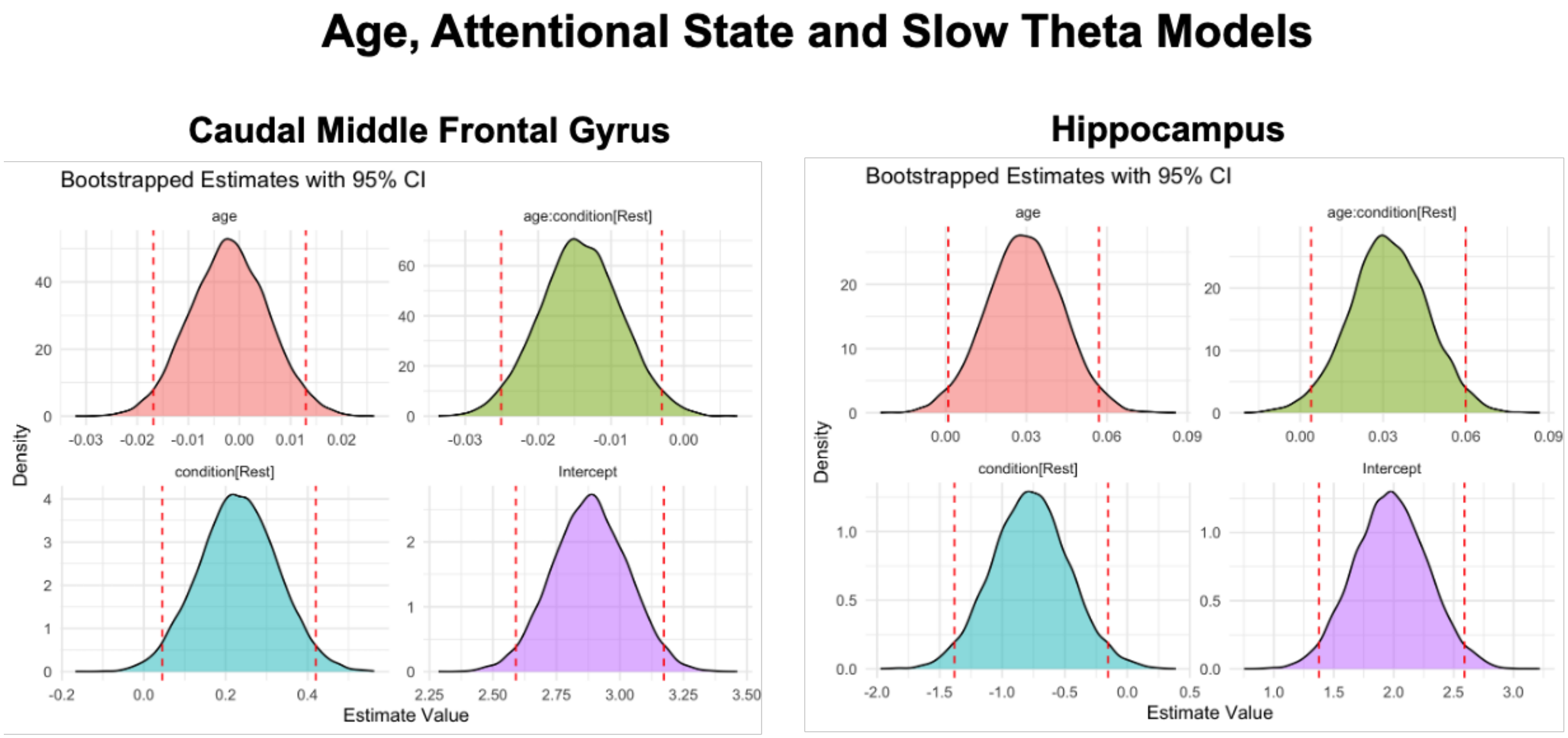


**S9. Bootstrapped beta coefficients for age, attentional state and slow theta models for the regions that survived multiple comparison correction (left, caudal middle frontal gyrus; right, hippocampus).** Bootstrapped confidence intervals for the intercept, main effects, and interactions. Y-axis represents the density of the bootstrapped coefficients, and the x-axis represents the estimated beta values. The dashed red lines indicate the 95% confidence interval from the original mixed-model estimates.

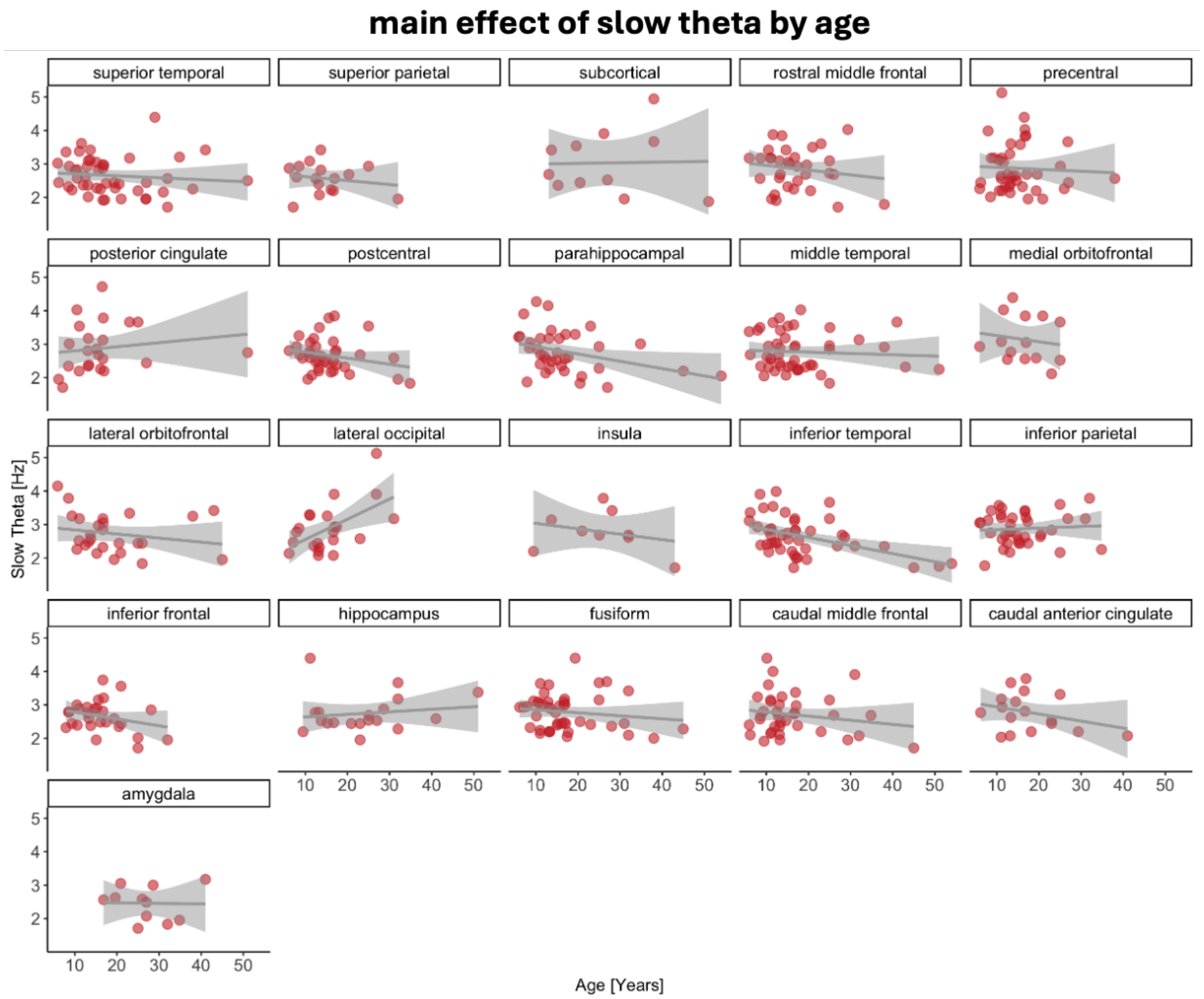


**S10. Relationship between the slow theta and age.** Peak slow theta frequencies are on the y-axis, with higher values denoting faster frequencies. Age is on the x-axis, with higher values denoting older age (in years). Data points indicate individual subjects, collapsed across channel and condition (task-based, task-free).

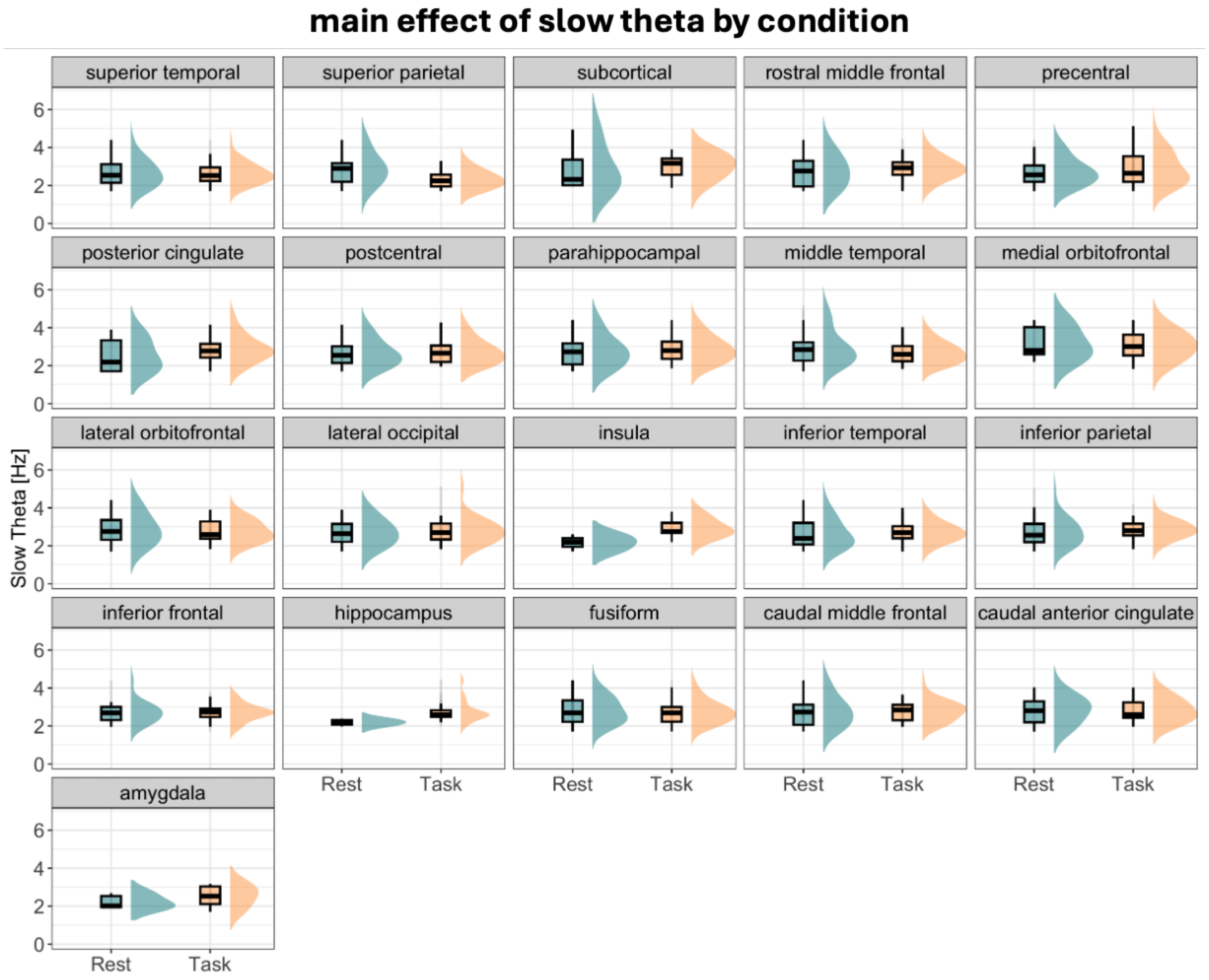


**S11. Differences in the peak slow theta frequencies between conditions (task-based, task-free/rest).** Peak slow theta frequencies are on the y-axis, with higher values denoting faster frequencies. Condition is on the x-axis.

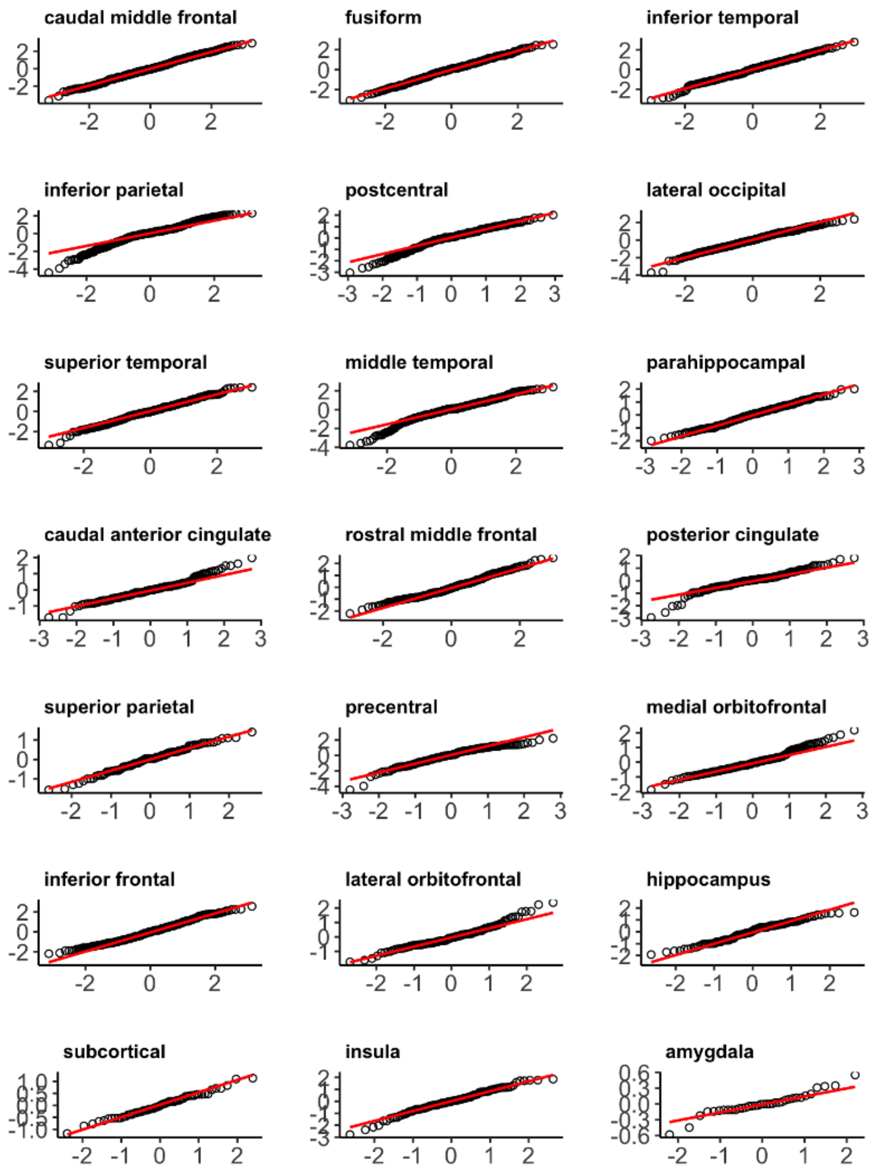


**S12.** Q-Q Plots of residuals for models examining age and attentional state on peak fast theta for each ROI.

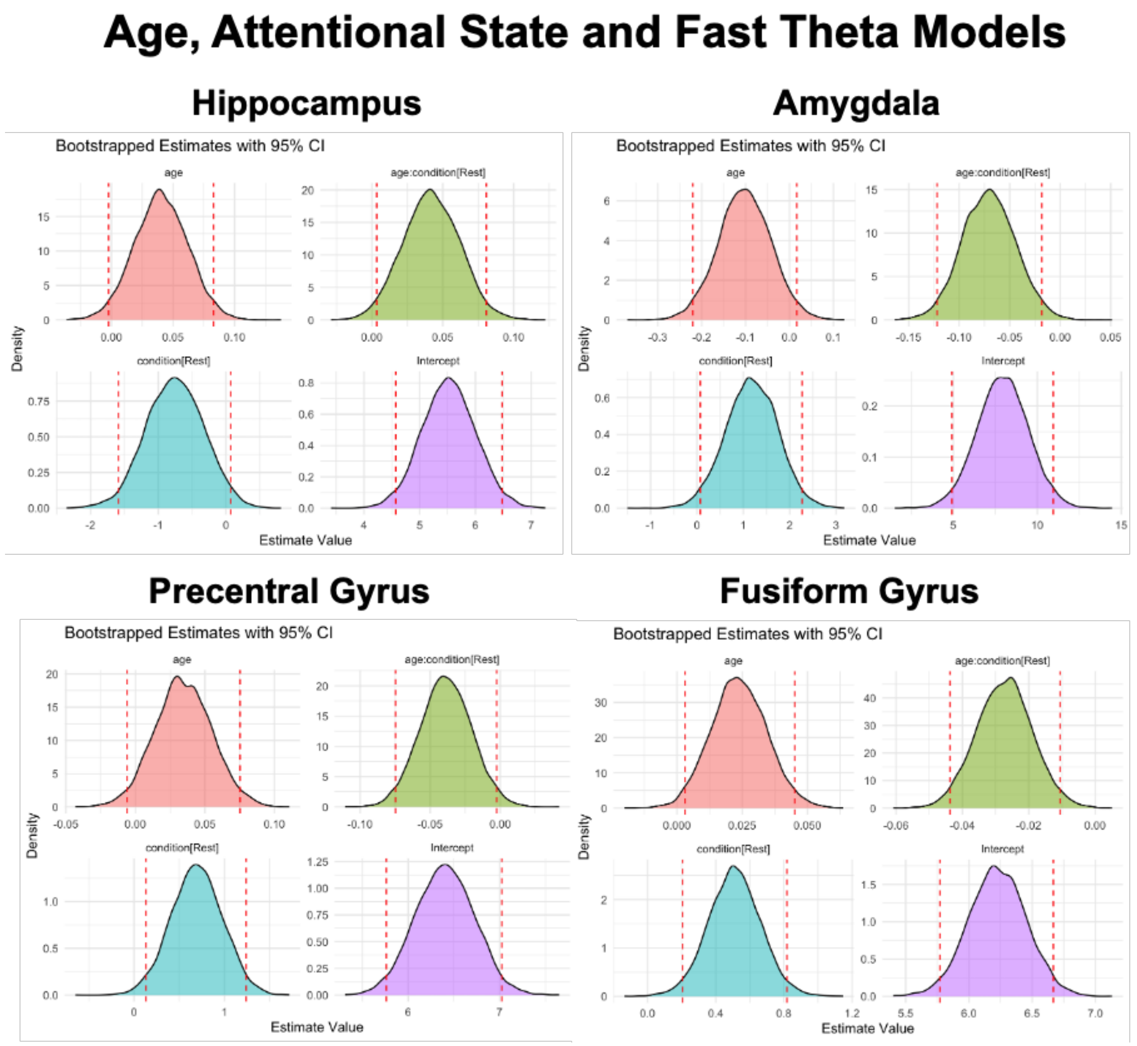


**S13. Bootstrapped beta coefficients for age, attentional state and fast theta models for the regions that survived multiple comparison correction (top left, hippocampus; top right, amygdala; bottom left, precentral gyrus; bottom right, fusiform gyrus).** Bootstrapped confidence intervals for the intercept, main effects, and interactions. Y-axis represents the density of the bootstrapped coefficients, and the x-axis represents the estimated beta values. The dashed red lines indicate the 95% confidence interval from the original mixed-model estimates.

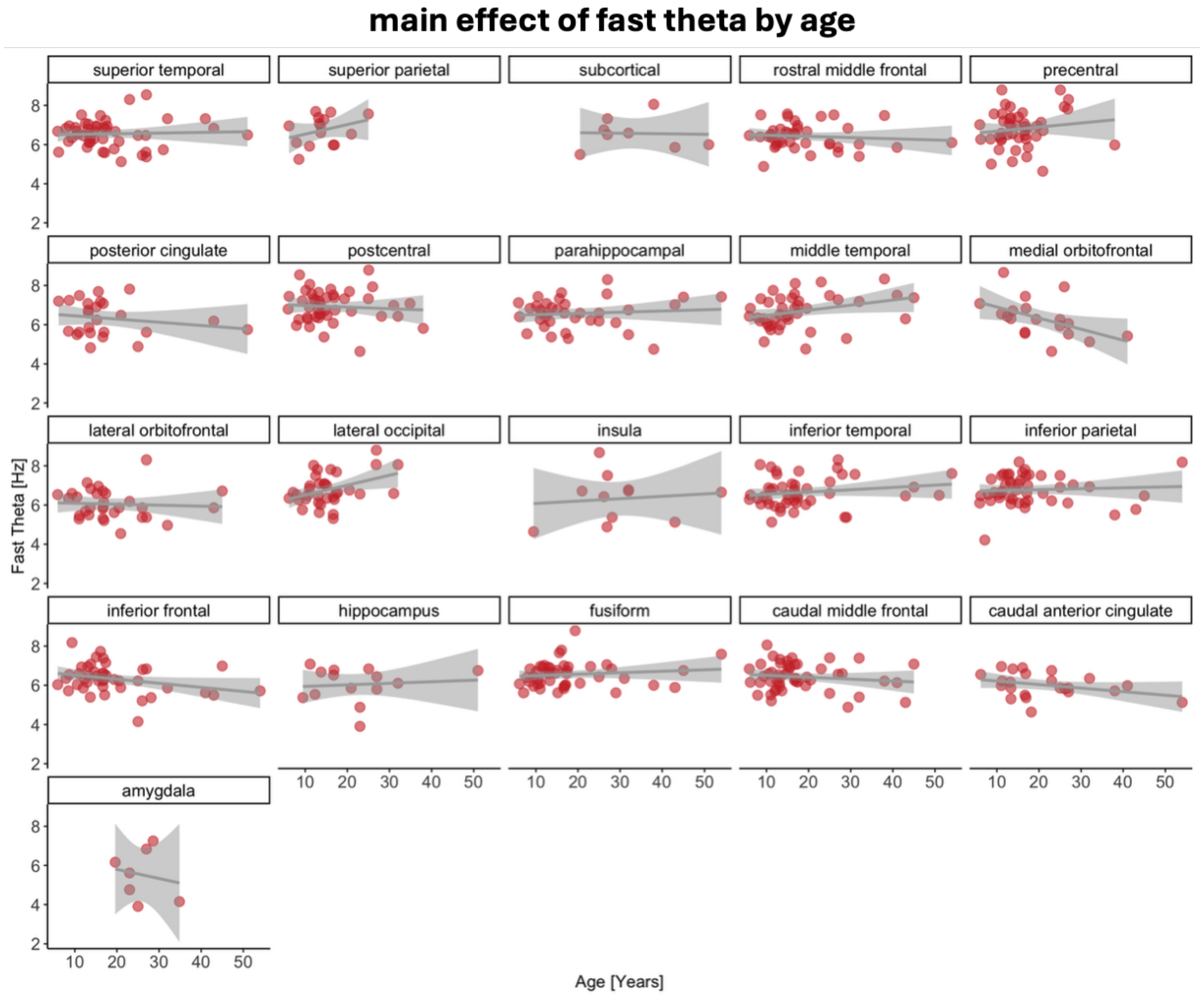


**S14. Relationship between the fast theta and age.** Peak fast theta frequencies are on the y-axis, with higher values denoting faster frequencies. Age is on the x-axis, with higher values denoting older age (in years). Data points indicate individual subjects, collapsed across channel and condition (task-based, task-free).

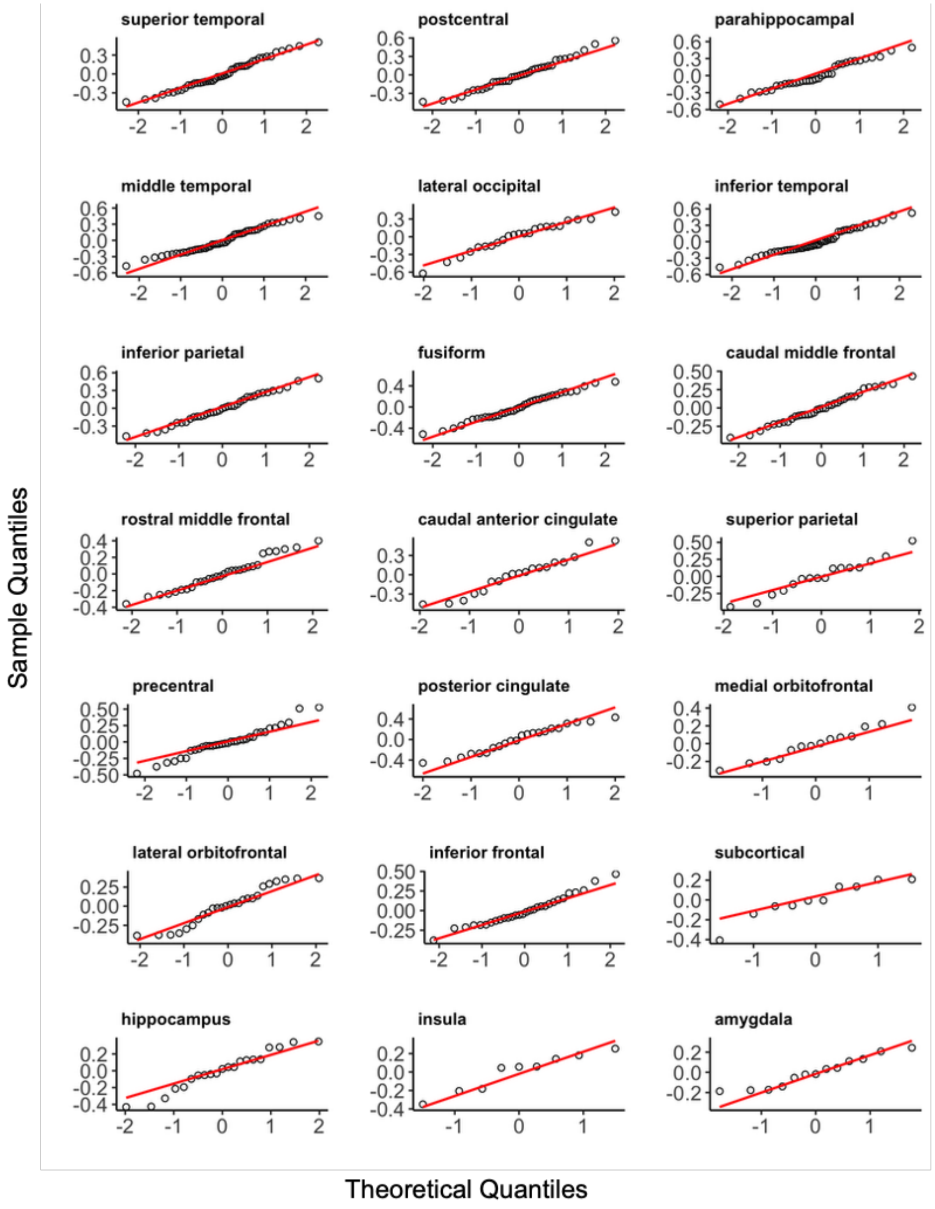


**S15.** Q-Q Plots of residuals for models examining task-based slow theta on memory for each ROI.

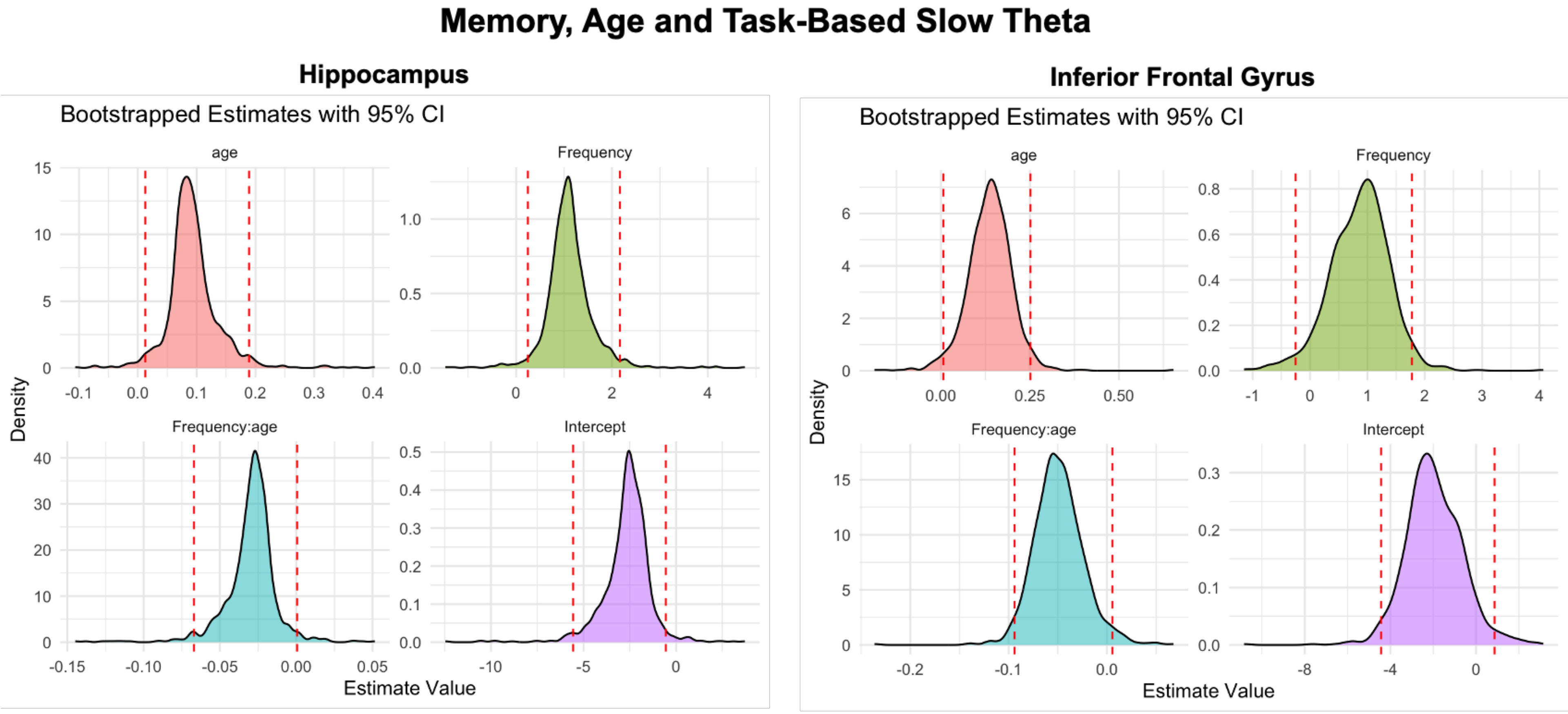


**S16. Bootstrapped beta coefficients for age, and task-based slow theta on memory for the regions that survived multiple comparison correction (left, hippocampus; right, inferior frontal gyrus).** Bootstrapped confidence intervals for the intercept, main effects, and interactions. Y-axis represents the density of the bootstrapped coefficients, and the x-axis represents the estimated beta values. The dashed red lines indicate the 95% confidence interval from the original mixed-model estimates.

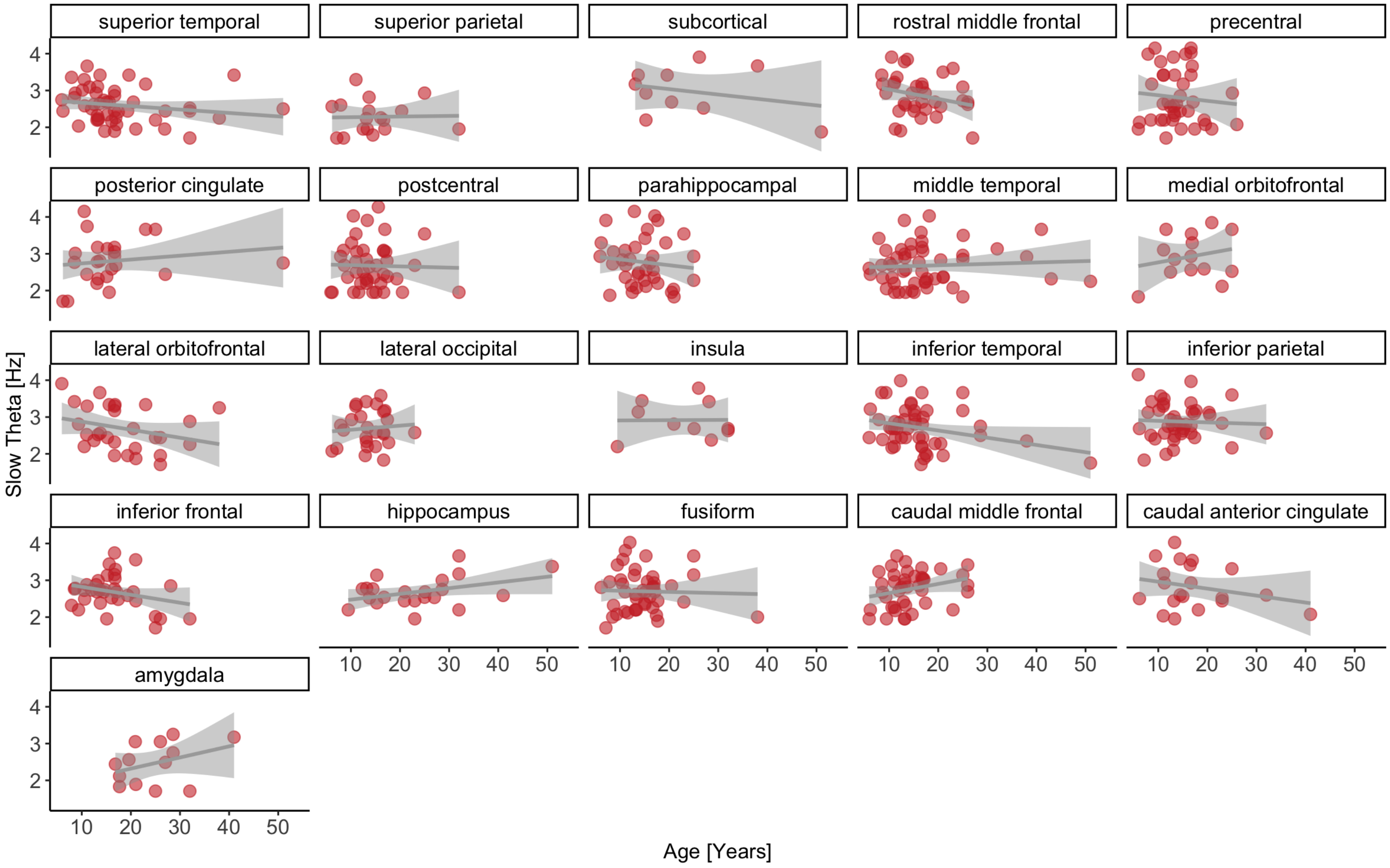


**S17. Relationship between task-based slow theta on memory.** Peak slow theta frequencies are on the y-axis, with higher values denoting fast frequencies. Age is on the x-axis, with higher values denoting older age (in years). Data points indicate individual subjects, collapsed across channel.

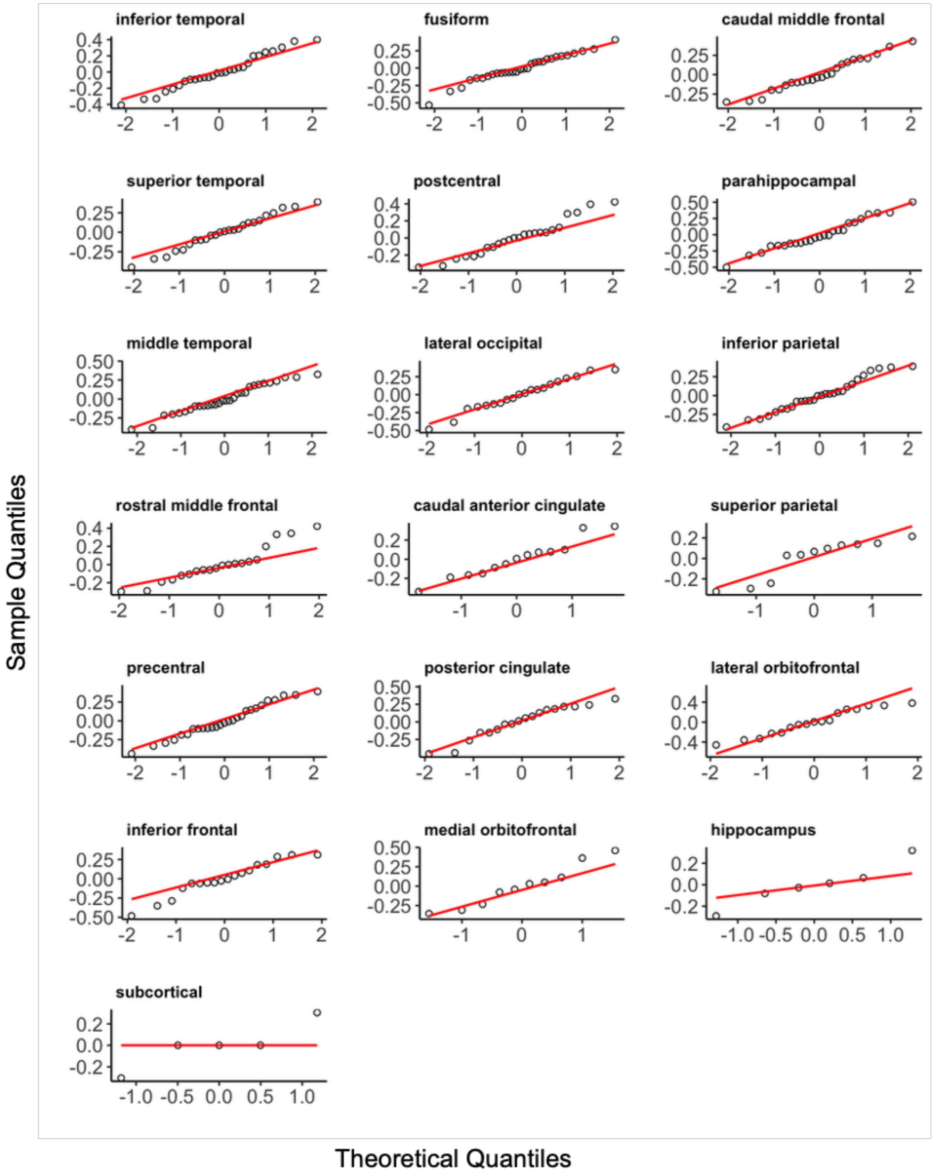


**S18.** Q-Q Plots of residuals for models examining task-free slow theta on memory for each ROI.

#### Memory, Age and Task-Free Slow Theta

##### Fusiform Gyrus

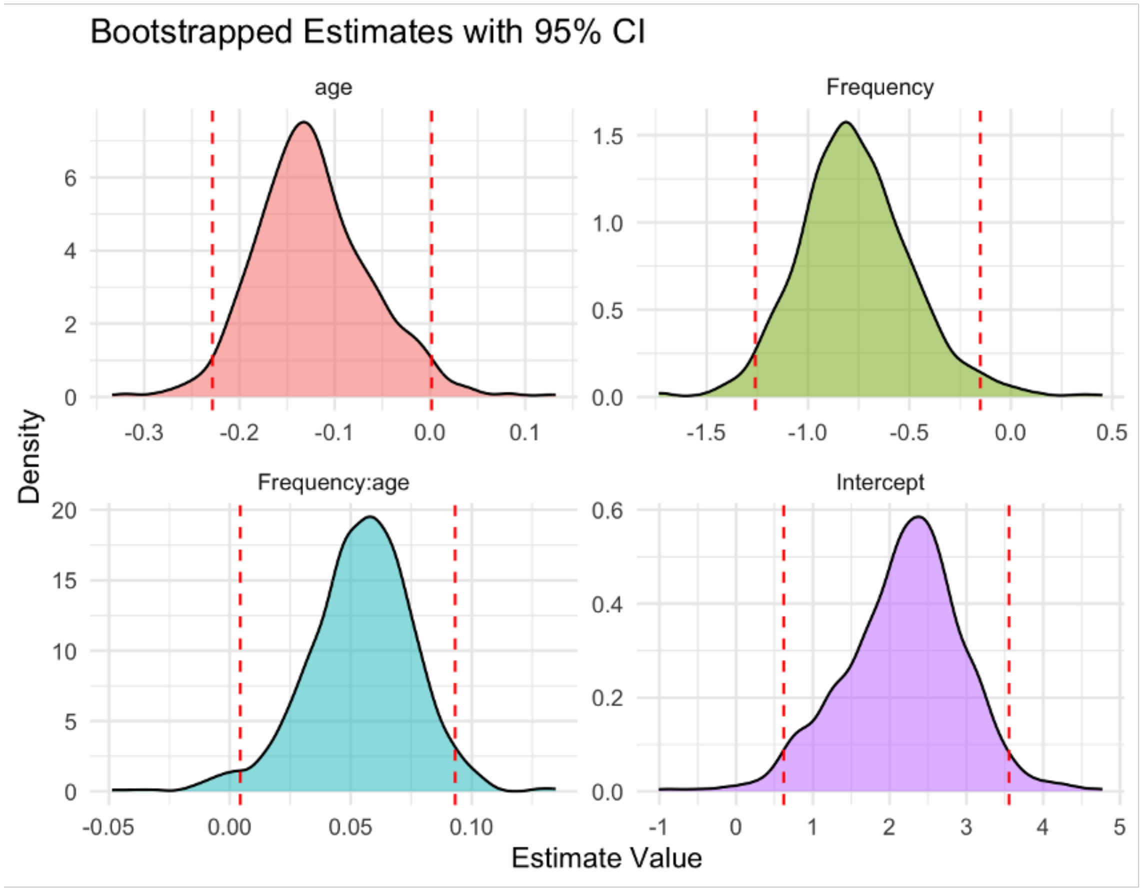


**S19. Bootstrapped beta coefficients for age, and task-free slow theta on memory for the regions that survived multiple comparison correction (fusiform gyrus).** Bootstrapped confidence intervals for the intercept, main effects, and interactions. Y-axis represents the density of the bootstrapped coefficients, and the x-axis represents the estimated beta values. The dashed red lines indicate the 95% confidence interval from the original mixed-model estimates.

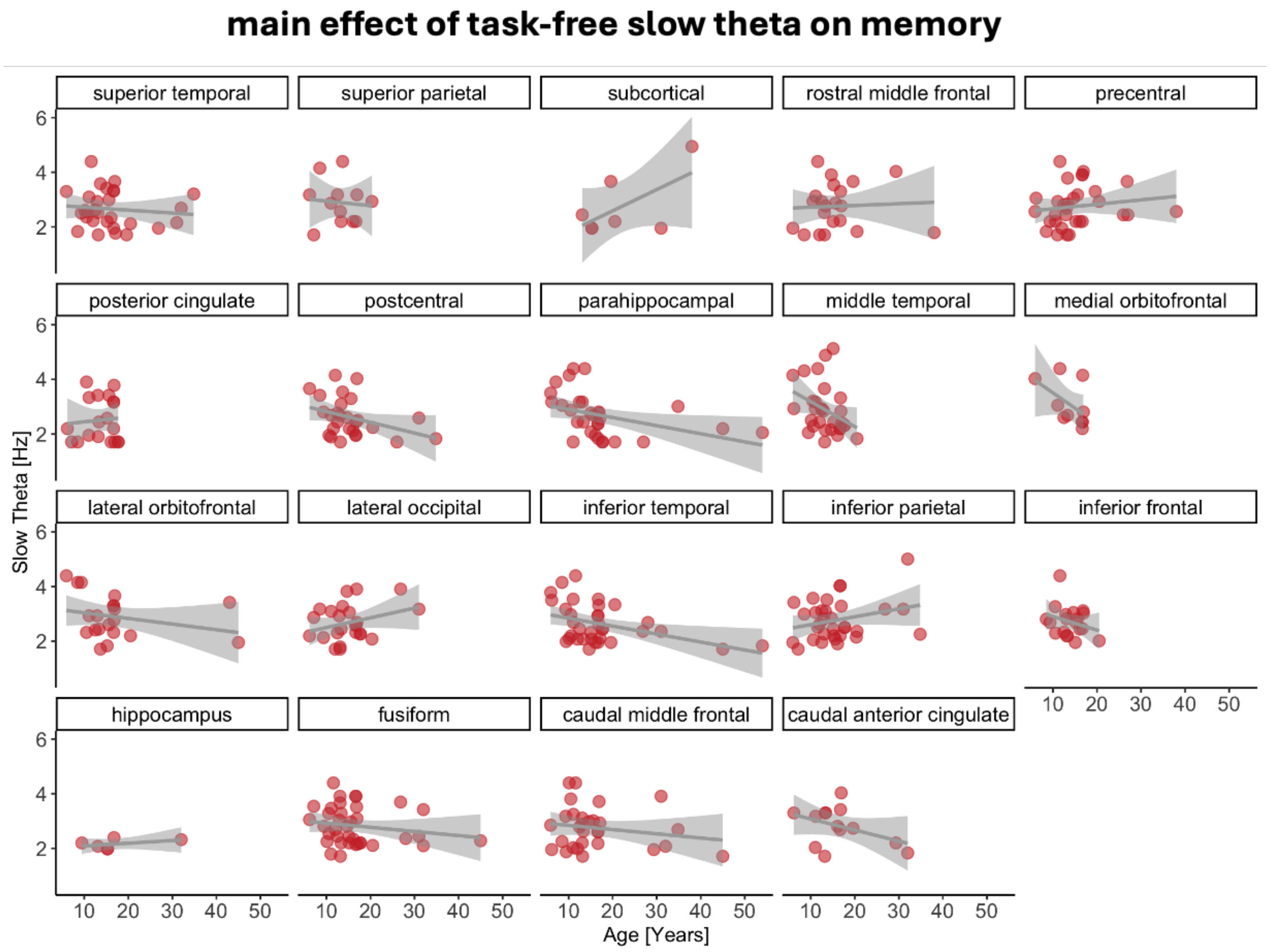


**S20. Relationship between task-free slow theta on memory.** Peak slow theta frequencies are on the y-axis, with higher values denoting fast frequencies. Age is on the x-axis, with higher values denoting older age (in years). Data points indicate individual subjects, collapsed across channel.

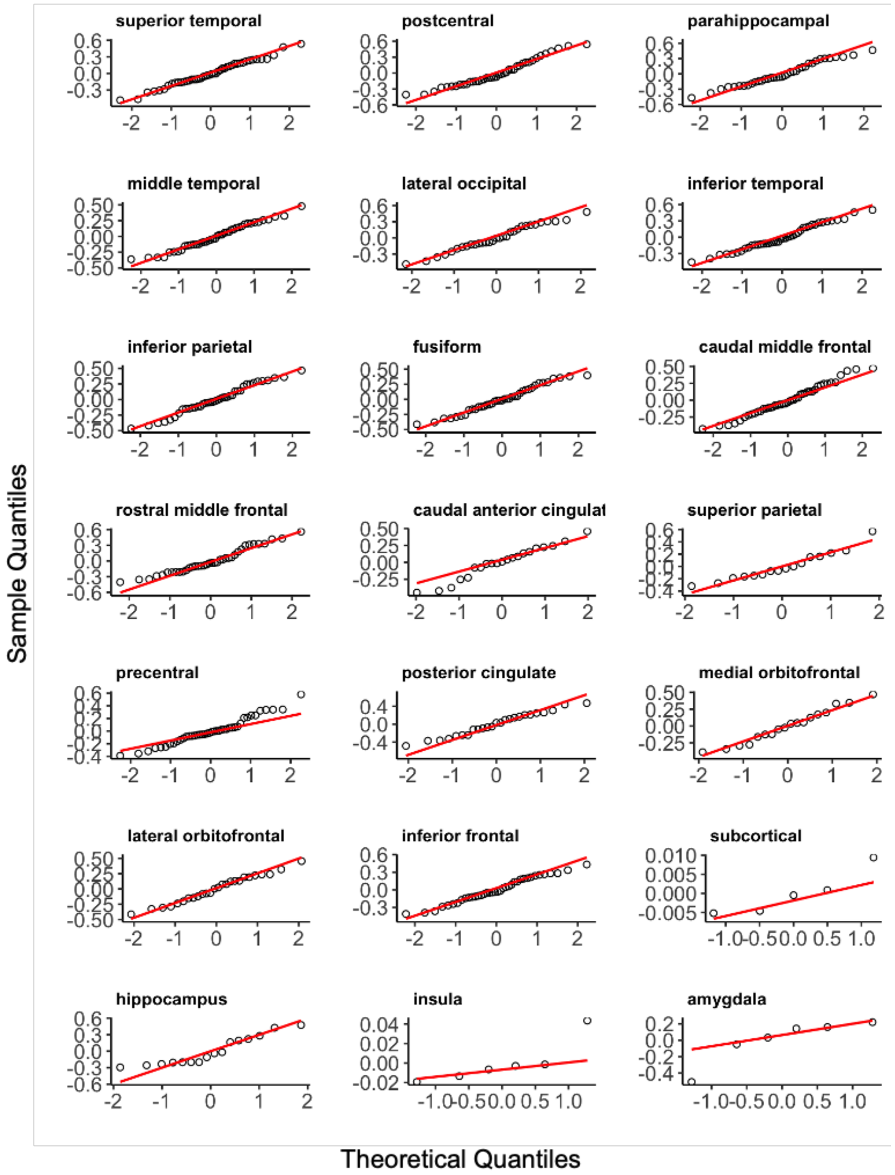


**S21.** Q-Q Plots of residuals for models examining task-based fast theta on memory for each ROI.

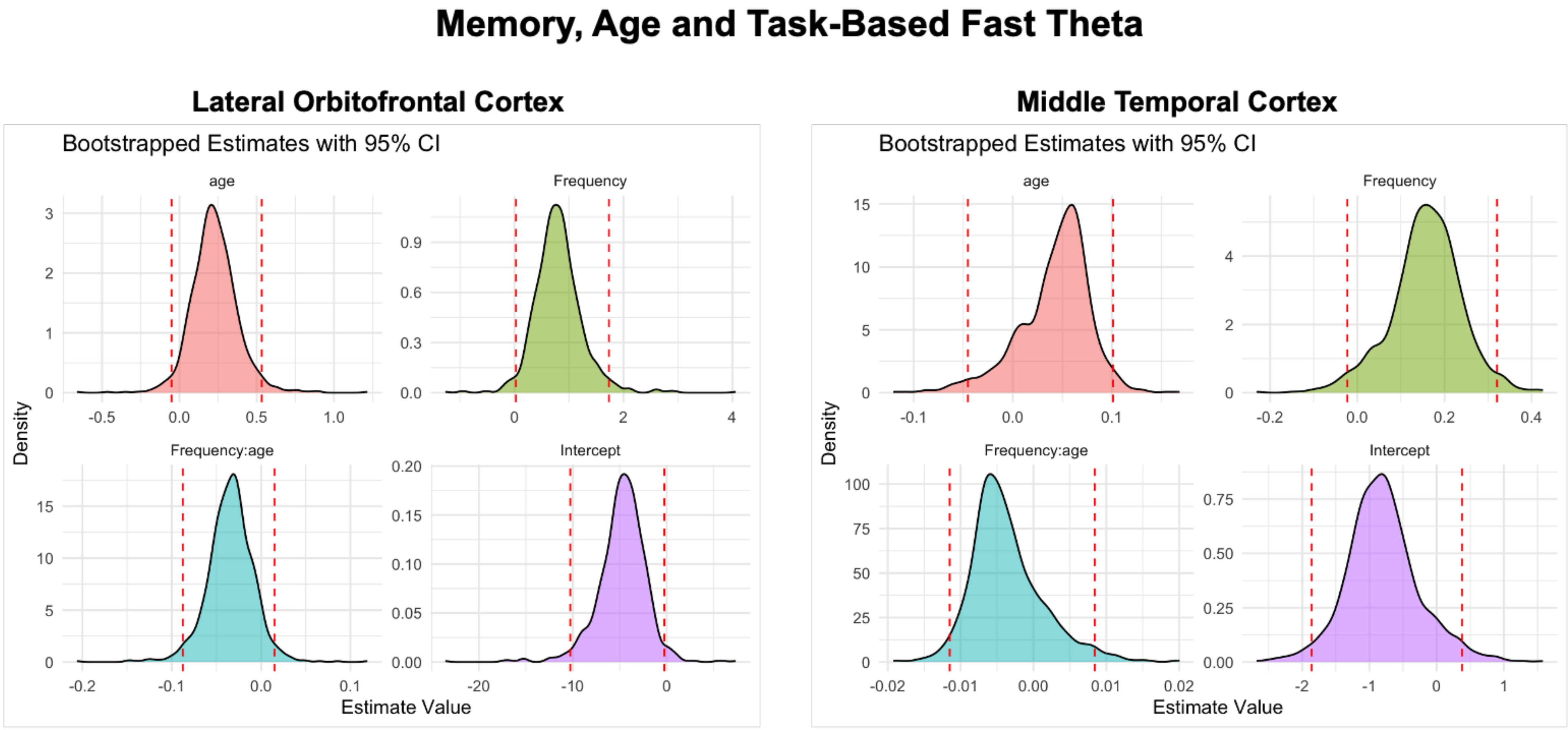


**S22. Bootstrapped beta coefficients for age, and task-based fast theta on memory for the regions that survived multiple comparison correction (left, lateral orbitofrontal cortex; right, middle temporal cortex).** Bootstrapped confidence intervals for the intercept, main effects, and interactions. Y-axis represents the density of the bootstrapped coefficients, and the x-axis represents the estimated beta values. The dashed red lines indicate the 95% confidence interval from the original mixed-model estimates.

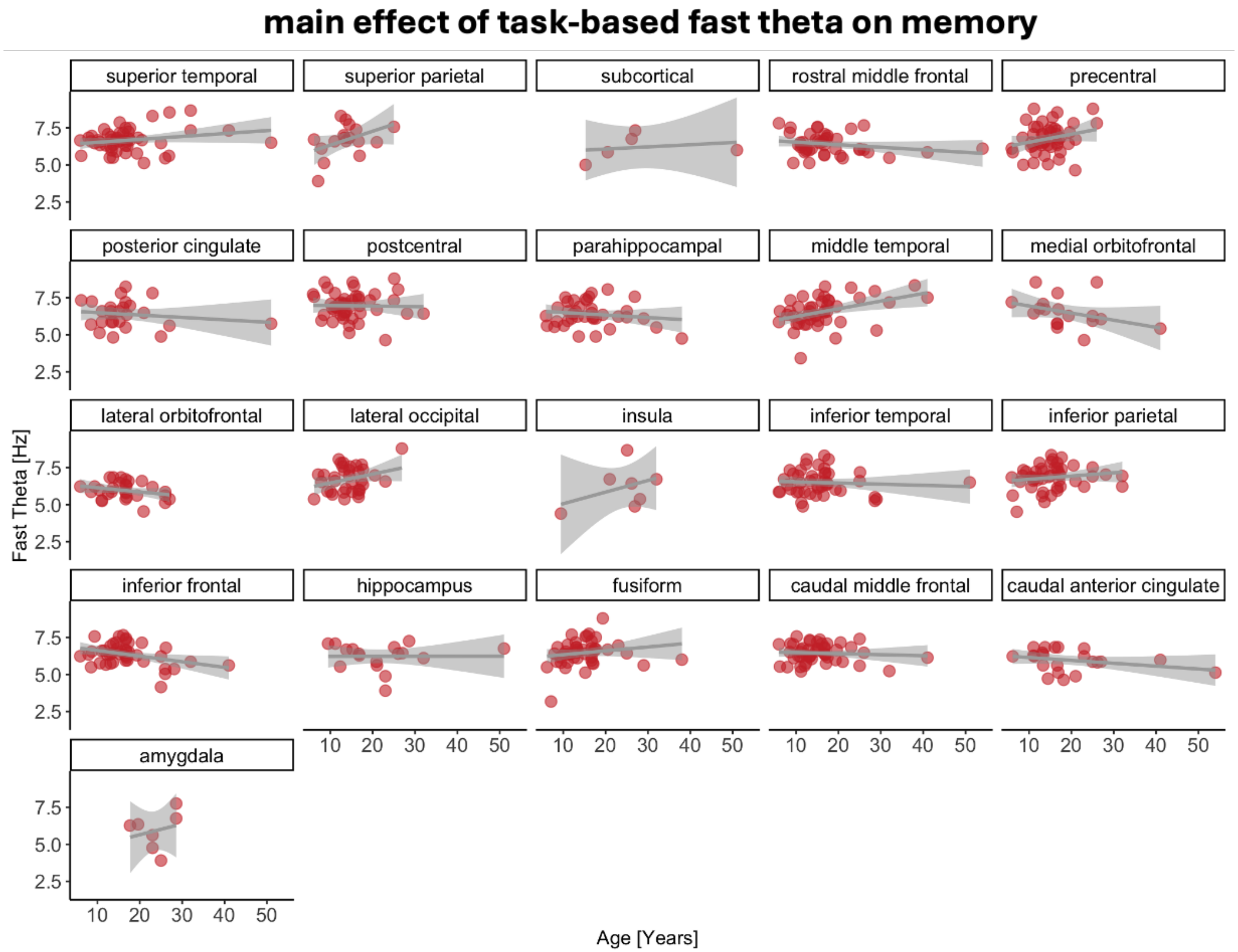


**S23. Relationship between task-based fast theta on memory.** Peak fast theta frequencies are on the y-axis, with higher values denoting fast frequencies. Age is on the x-axis, with higher values denoting older age (in years). Data points indicate individual subjects, collapsed across channel.

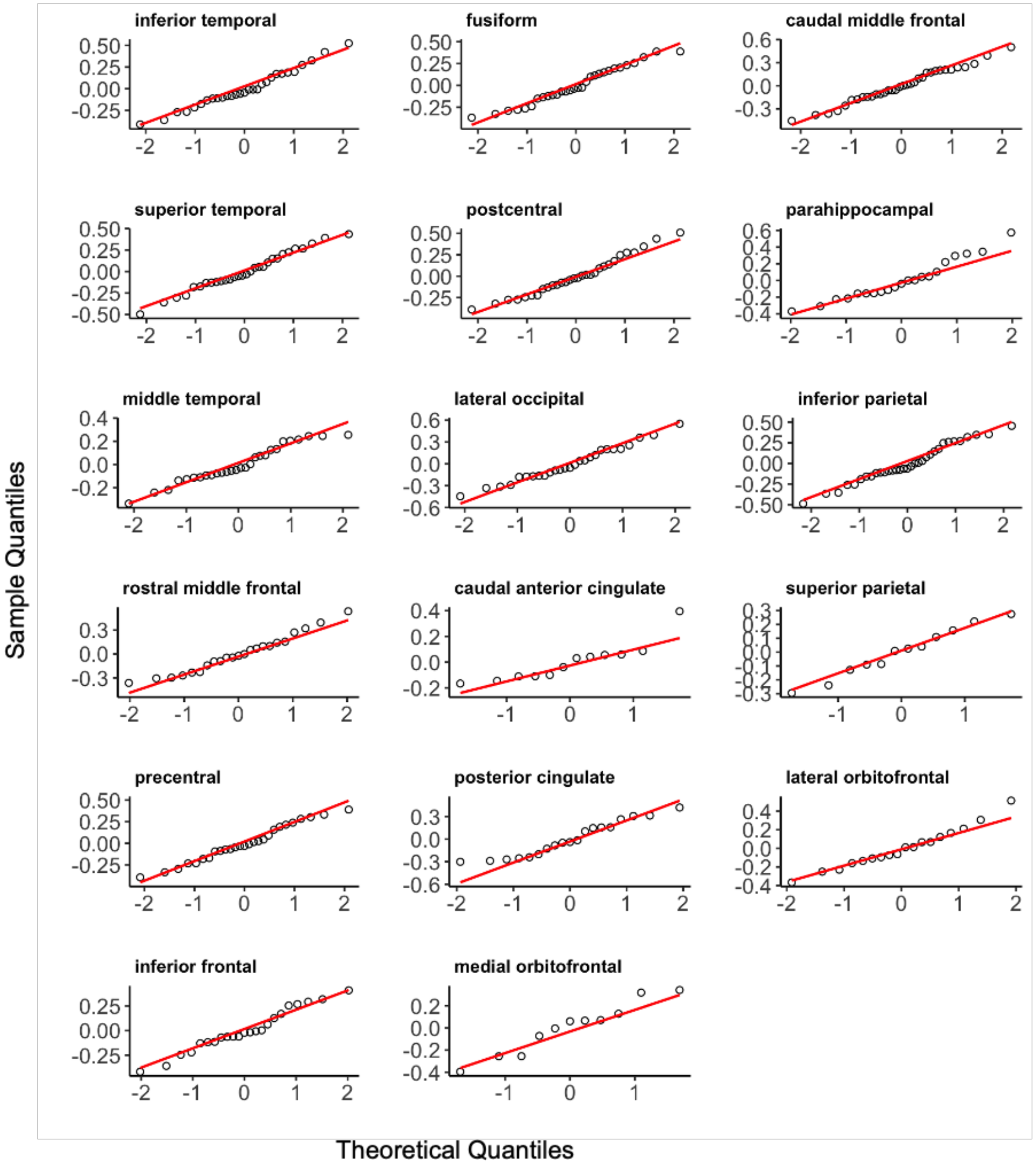


**S24.** Q-Q Plots of residuals for models examining task-free fast theta on memory for each ROI.

### Memory, Age and Task-Free Fast Theta

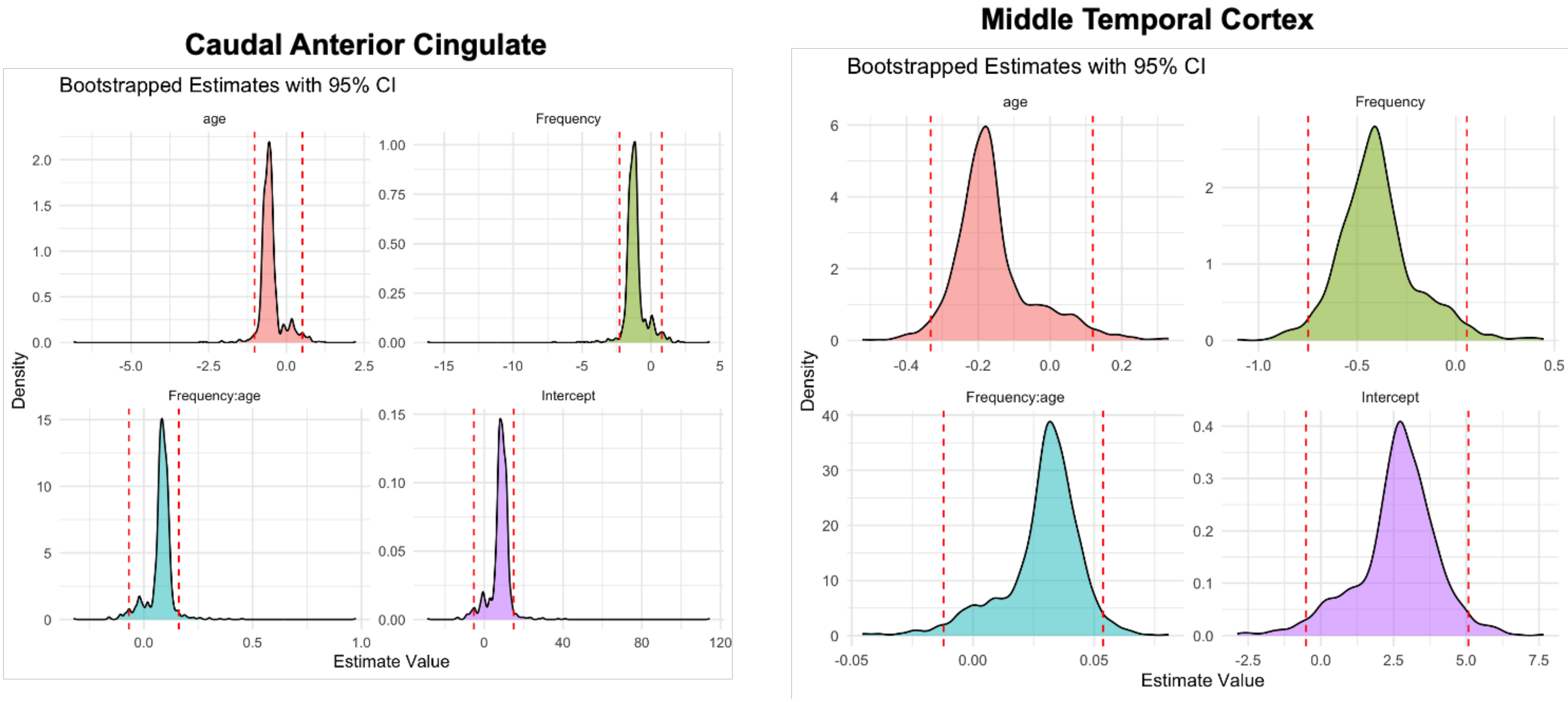


**S25. Bootstrapped beta coefficients for age, and task-free fast theta on memory for the regions that survived multiple comparison correction (left, caudal anterior cingulate; right, middle temporal cortex).** Bootstrapped confidence intervals for the intercept, main effects, and interactions. Y-axis represents the density of the bootstrapped coefficients, and the x-axis represents the estimated beta values. The dashed red lines indicate the 95% confidence interval from the original mixed-model estimates.

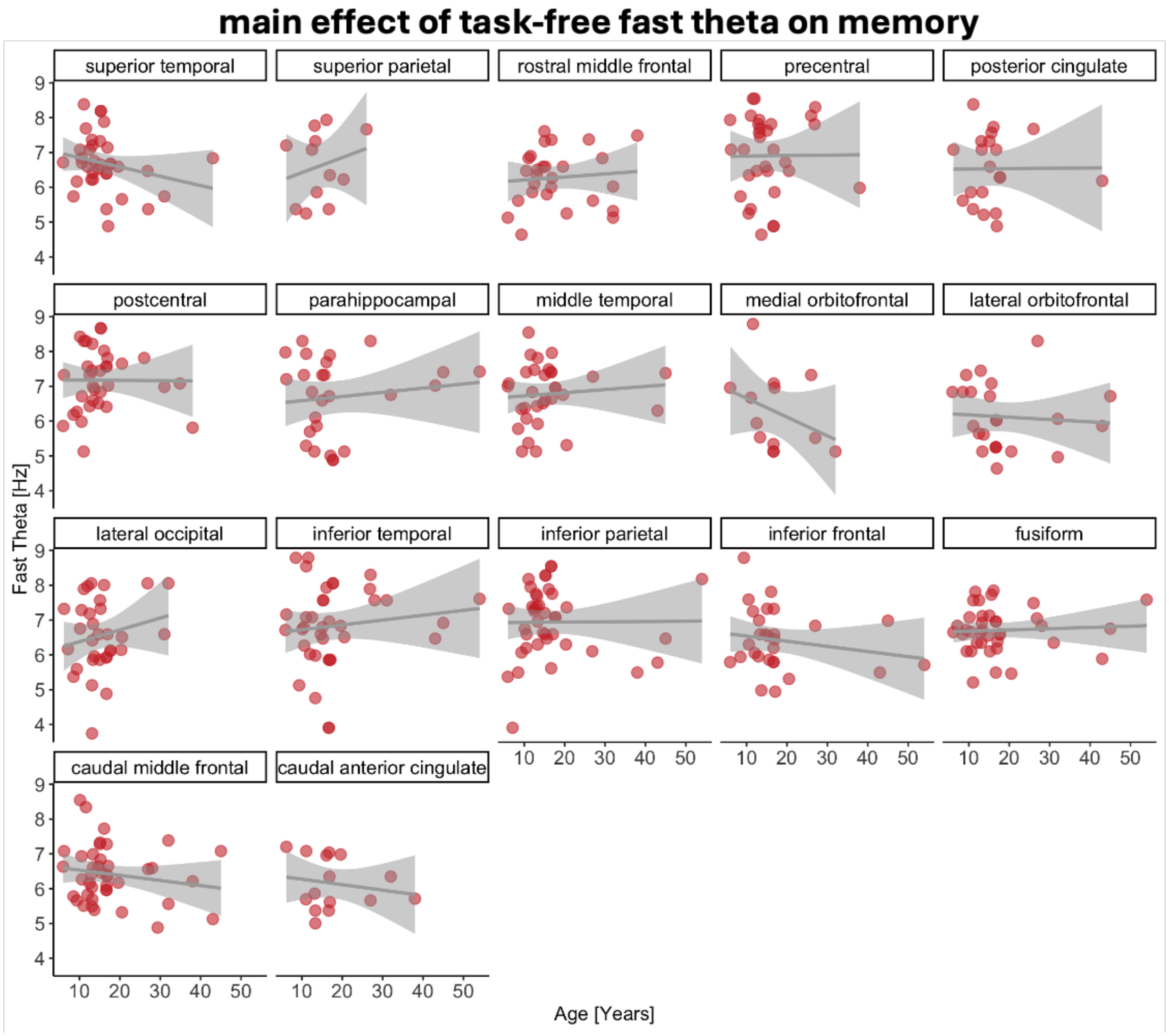


**Figure S26. Relationship between task-free fast theta on memory.** Peak fast theta frequencies are on the y-axis, with higher values denoting fast frequencies. Age is on the x-axis, with higher values denoting older age (in years). Data points indicate individual subjects, collapsed across channel.

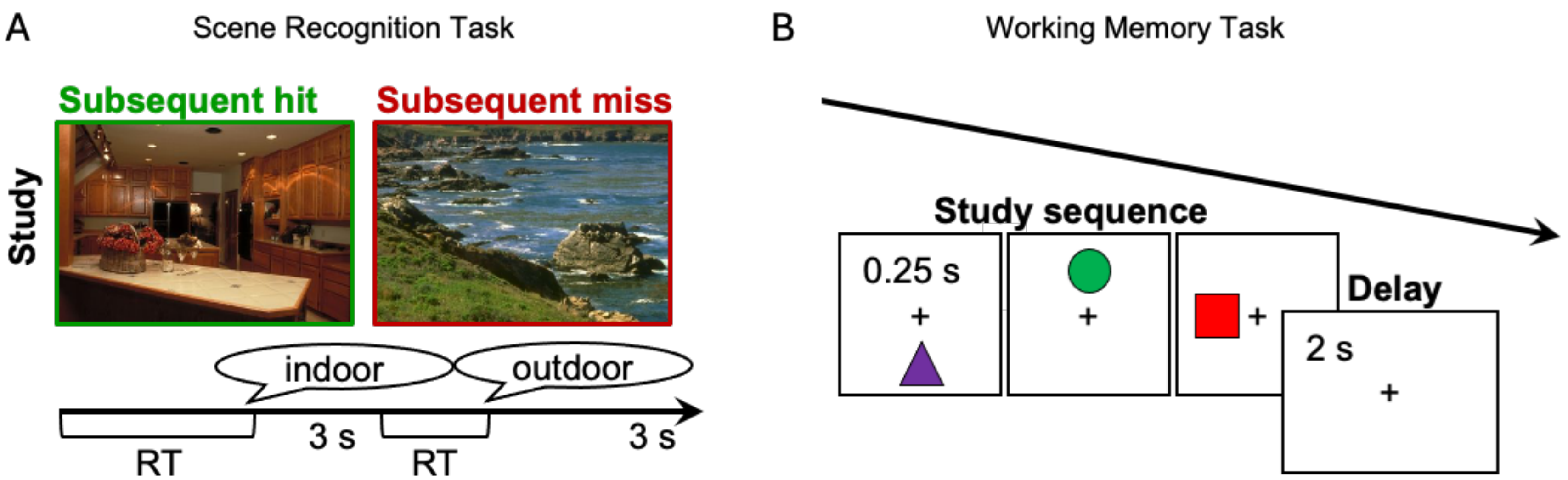


**S27. Schematic of the two visual recognition memory tasks. (A).** Illustration of the visual scene recognition task. Subjects studied sets of 40 pictures of scenes (3s each, separated by a 500ms interstimulus fixation) and made an indoor/outdoor judgment of each scene in preparation for a recognition memory test of all scenes presented during the study block, intermixed with 20 new scenes. **(B)** Illustration of the visual working memory task. Subjects encoded three shapes in a specific spatiotemporal sequence in preparation for a self-paced old/new recognition test of sequences that match exactly or mismatch on one dimension (i.e., shape identity, spatial position, or temporal order).
